# Oral immune priming modulates microbiota composition and supports pathogen control in the Manila clam (*Ruditapes philippinarum*)

**DOI:** 10.64898/2025.12.27.696656

**Authors:** Bruno K. Rodino-Janeiro, Diego Rey-Varela, Ana L. Diéguez, Sergio Rodriguez, Clara Martinez, Javier Dubert

## Abstract

Immunological memory was long considered an exclusive feature of vertebrates. However, extensive evidence now shows that invertebrates possess forms of innate immune memory—known as immune priming— where previous exposure to a pathogen enhances subsequent immune responses and host protection. Immune priming has been proposed as a promising strategy for disease prevention in shellfish aquaculture. However, immune priming remains largely unexplored in marine bivalves, particularly in the Manila clam (*Ruditapes philippinarum*), a top-ten global aquaculture species. Here, we investigated for the first time the effects of oral immune priming on host survival, pathogen dynamics, and microbiota composition in *R. philippinarum* against the emergent bivalve pathogen *V. europaeus*. Priming was induced using the live bacterial pathogen at a sublethal dose, followed by a lethal secondary exposure. Primed clams exhibited a significant survival following the second challenge (87% survival vs. 0% in non-primed clams), demonstrating robust protection against reinfection. Quantitative PCR (qPCR) revealed that primed clams rapidly reduced pathogen loads after 48 h during the second challenge, reaching concentrations below the mortality threshold observed in non-primed clams (∼10^5^ copies mg⁻¹). Interestingly, the pathogen was able to persist at low and non-harmful concentration (∼10^2^ copies mg⁻¹) in primed clams along both challenges. Full-length 16S rRNA metabarcoding analyses showed that immune priming shifts the host microbiota. Alpha and beta diversity indicated a progressive reduction in diversity and the establishment of a specific and resilient bacterial community in primed clams. Clustering analyses identified a priming-associated microbiota dominated by *Acinetobacter*, *Brevundimonas*, *Sphingobium*, and *Psychrobacter*, which persisted through the secondary challenge but was absent or depleted in non-primed clams. Conversely, members of the *Arcobacteraceae* (e.g., *Arcobacter*, *Poseidonibacter*) were absent after priming and emerged only during second infection, decreasing in primed clams but increasing in non-primed clams coinciding with high mortalities. Our findings provide the first phenotypic and microbiome-level evidence of oral immune priming in Manila clam. Here we demonstrate that priming enhances pathogen control and promotes the establishment of a protective microbiota that may interact with the host immune system to confer resistance against bacterial infection. These results open new avenues for immune-priming and microbiota-based strategies to improve disease resistance in bivalve aquaculture.

## 1 Introduction

Training immune memory through vaccination is one of the most powerful strategies against infectious diseases. Immunological memory was considered for decades an exclusive trait for vertebrates, standing out jawed vertebrates as the most developed adaptive immune system in Chordata class. This feature is linked to the adaptive immune system mediating the response to exogenous antigens through the production of specific antibodies using somatic rearrangement and clonal expansion of lymphocytes (Sompayrac, 2022). This allows individuals to acquire immunity to emerging pathogens, rapidly adapting to new challenges. However, over the past two decades, several studies have revealed the existence of an antigen-independent immunological memory in invertebrates (Pradeu & Du Pasquier, 2018). This trait is termed as innate immune memory, trained immunity, or immune priming (Low & Chong, 2020; Montagnani et al., 2024). Immune priming triggers an effect similar to vaccination, where an exposure to a pathogen, using live—at sub-lethal doses— or inactivated microbes—by chemical or physical methods (e.g., heat-shock, formalin…)—can enhance immune responses and confer protection against subsequent infections by the same or even unrelated pathogens (Lanz-Mendoza & Contreras-Garduño, 2022; Low & Chong, 2020; Melillo et al., 2018; Montagnani et al., 2024; Pradeu & Du Pasquier, 2018). Despite invertebrates comprising more than 95% of animal diversity, immune priming has been demonstrated in only a few taxa, including cnidarians, molluscs, and several ecdysozoan orders (Lanz-Mendoza & Contreras-Garduño, 2022; Low & Chong, 2020; Melillo et al., 2018; Montagnani et al., 2024; Pradeu & Du Pasquier, 2018). Due to the high variability of these kinds of systems, the underlying mechanisms driving to the immune priming cannot be translated between species as different phyla respond to the immunizing stimulus differently (Lanz-Mendoza & Contreras-Garduño, 2022).

In marine bivalves, evidence for immune priming remains limited but highly promising (Montagnani et al., 2024). This invertebrate group is relevant as they are among the most economically and environmentally sustainable aquaculture species, requiring no external feeding inputs and providing protein of high nutritional value and important socio-economic benefits to coastal communities (FAO, 2024; Naylor et al., 2021). Globally, Pacific oyster (*Magallana gigas*) and Manila clam (*Ruditapes philippinarum*) account for over 8% of total aquaculture production (FAO, 2024). However, microbial diseases have become one of the main constraints for global bivalve aquaculture. These diseases, caused primarily by bacteria, viruses, and protozoan parasites, are responsible for recurrent mass mortalities that threaten the sustainability and economic viability of this sector (Dubert et al., 2017a). Bivalves’ filter-feeding behaviour makes them particularly vulnerable: they bioaccumulate pathogens from the surrounding water/animals and can release them back into the environment, facilitating rapid disease spread (Zannella et al., 2017). The increasing frequency and severity of these microbial diseases—exacerbated by climate change and by aquaculture-specific factors such as high stocking densities or the increase of imports/exports — affecting both wild and farmed marine bivalves pose growing challenges for aquaculture sustainability (Dubert et al., 2017a; Naylor et al., 2021; Smolowitz, 2024).

Among bacterial diseases, vibriosis caused by pathogenic *Vibrio* spp. (*V. europaeus, V. coralliilyticus*, among others) is a major threat to bivalve production (Dubert et al., 2017a). Despite over five decades of research, no effective and eco-friendly prophylactic or therapeutic treatments exist to fight pathogenic vibrios in bivalve aquaculture (Dubert et al., 2017a; Smolowitz, 2024). For instance, in aquaculture settings such as hatcheries, antibiotics remain widely used in the culture tanks to protect bivalves in early life stages (larvae and seed). However, their misuse has accelerated the emergence and horizontal transfer of antimicrobial resistance (AMR) genes among *Vibrio* spp. in these facilities (Dubert et al., 2017a; Naylor et al., 2021). Moreover, delivering preventive treatments to bivalves in open marine environment is technically difficult and often impracticable. At this point, immune priming appears as a promising strategy for addressing this gap in bivalve aquaculture providing a long-lasting protection against pathogens (Montagnani et al., 2024). Experimental evidences of immune priming include bivalve species with interest in aquaculture such as the Japanese scallop (*Chlamys farreri*)(Cong et al., 2008, 2009; Wang et al., 2013; Yue et al., 2013), mussels (*Mytilus galloprovincialis* (Rey-Campos et al., 2019) and *Perna viridis* (Aleng et al., 2015), and the Pacific oyster, where immune priming has been studied extensively against both bacterial and viral pathogens (Gerdol & Venier, 2015; Green et al., 2016; Lafont et al., 2017, 2020; T. Zhang et al., 2014). Despite being one of the most important species in global aquaculture (FAO, 2024), the Manila clam has been largely overlooked in immune priming research. To date, only one recent study demonstrated this phenomenon in this species, and it focused exclusively on the host immune response to *V. anguillarum* infection (Li et al., 2025).

Among *Vibrio* pathogens, *Vibrio europaeus* has emerged over the last decade as a particularly relevant threat, being responsible for recurrent and massive mortality events in shellfish hatcheries in France, Spain, Chile, and the United States—countries that rank among the world’s top ten bivalve producers (Dubert et al., 2017a; Rojas et al. 2021). This bacterial species exhibits a broad host range, infecting several key bivalve species such as the Manila clam, Pacific oyster, and Chilean scallop (*Argopecten purpuratus*), as well as gastropod mollusks such as abalones (*Haliotis* spp.) (Saulnier et al., 2010). Importantly, its virulence has been experimentally demonstrated across multiple life stages of bivalves, including larvae of oysters (*C. gigas* and *Ostrea edulis*), clams (*R. decussatus*, *R. philippinarum*, *Donax trunculus*, *Ensis arcuatus*, and *Polititapes romboides*), and scallops (*A. purpuratus*); post-larvae (or early spat) of Manila clams; and juveniles of Pacific oysters (Prado et al. 2005, 2015; Travers et al., 2014; Mersni-Achour et al., 2014, 2015; Dubert et al., 2016, 2017b; Rojas et al., 2021). Owing to its wide host spectrum and strong association with hatchery mortality outbreaks, *V. europaeus* constitutes a highly relevant bacterial model for investigating vibriosis in bivalves. These characteristics make it particularly suitable for exploring alternative disease mitigation strategies, including immune priming-based approaches aimed at enhancing host resistance to infection.

The interactions between the immune system, microbiota and pathogens, are too often studied separately rather than in a holistic way. When studying bipartite interactions such as host–microbiota relationships, numerous studies—mainly in insect models—have demonstrated that the microbiota plays a key role in activating immune priming. These studies demonstrate that symbiotic microorganisms can shape host immunity and contribute directly to immune priming by inducing immune effectors (e.g., AMPs) or modulating immune signaling pathways (e.g., the Toll pathway) (Futo et al., 2016; Hernández-Martínez et al., 2010; Horak et al., 2020; Jang et al., 2024; Keshavarz et al., 2025; Korša et al., 2022; Lang et al., 2022; Turner et al., 2024).

Despite the well-recognized importance of the immune-microbiota axis in insect models, the impact of immune priming on the microbiota of marine invertebrates remains poorly characterized and has so far been investigated exclusively in the Pacific oyster (*Magallana gigas*) (Fallet et al., 2022). Here, we present the first study to assess the effect of oral immune priming on the microbiota of the Manila clam against *V. europaeus*. This study aimed to deliver the first evidence of oral immune priming in the Manila clam, utilizing a novel and comprehensive approach for this bivalve species that integrates host survival, pathogen dynamics, and full-length 16S rRNA microbiome profiling. We then examined shifts in the host-associated microbiota during the priming process, with the aim of evaluating bacterial diversity and identifying the predominant taxa shaping community composition. This approach is essential for advancing our understanding of the mechanisms underlying immune priming in marine invertebrates.

## 2 Materials and methods

### 2.1 Model organisms: Manila clam (*Ruditapes philippinarum*) and *Vibrio europaeus*

Experimental challenges were conducted using Manila clam juveniles (*Ruditapes philippinarum*, 13 ± 1 mm shell length, 8 months old). A total of 1,000 clams were supplied by Proameixa (Galicia, Spain) and acclimated prior to the experiment in aquaria at the University of Santiago de Compostela (USC). Clams were maintained until use at room temperature (RT; approximately 19 ± 1 °C) under aeration and regularly fed with EasyBooster25 (Easyreefs, Spain) following the manufacturer’s recommendations.

*Vibrio europaeus* strain CECT 8136^T^ was used to evaluate immune priming in Manila clams. This bacteria was routinely cultured on Tryptone–Soy agar or broth (TSA–2/TSB–2, Condalab) supplemented with 2% (w/v) NaCl at 25 °C for 24 h. For the challenges described below, bacterial suspensions were prepared in sterile seawater (SSW) from overnight cultures and adjusted to an optical density at 600 nm (OD_600_) of 1.0, corresponding to approximately 10^8^ colony forming units (CFU) mL^−1^. Serial tenfold dilutions were performed (i) to confirm bacterial counts by spreading on TSA–2 plates and the concentration is expressed as CFU mL^−1^, and (ii) to adjust the bacterial inoculum to the desired final concentration according to the objectives of each challenge assay.

### 2.2 Challenges assays

The experimental design consisted of two consecutive exposures of Manila clam juveniles to the pathogen (*V. europaeus*), oral priming and second challenge (Figure 1). Six tanks were used per condition—three for DNA sampling and three for monitoring survival rates—as follows:

(i) Non-primed clams (n=6 tanks) but infected with the pathogen during the second challenge, used as positive controls (NP, C+).
(ii) Primed clams infected during the second challenge (P+SC) (n=6 tanks) to evaluate increased survival associated with the immune priming response.
(iii) Primed tanks not infected during the second challenge (n=6 tanks), used as negative controls (P, C–) to discard potential negative effects of priming on host survival throughout the experimental period.

Briefly, the experimental design included two main steps:

(a) **Oral priming:** NP (C+) tanks (15 juveniles per tank) containing only 100 mL of filtered seawater (FSW; 0.22-µm Nalgene Rapid-Flow, Thermo Fisher). On the other hand, Manila clam juveniles from P (C-) and P+SC tanks (15 juveniles per tank) were orally primed by immersion in FSW containing *V. europaeus* at a sub-lethal dose (final concentration ∼10^6^ CFU mL^−1^). All tanks were kept with aeration for 7 days (168 hours post-priming, hpp), with food supply and water renewal at day 5.
(b) **Second challenge:** After one week, clams from both NP (C+) and P+SC tanks were exposed to a lethal dose of *V. europaeus* CECT 8136^T^ (final concentration ∼10^7^ CFU mL^−1^) to evaluate the immune priming response, following the infection protocol described by Martínez et al. (2022). P (C–) tanks were not infected and served as negative controls as described above. According to Martinez et al. (2022), infection protocol involves two steps: (i) infection: juveniles were transferred to infection tanks and immersed in the bacterial suspension for 24 h at room temperature (RT) without aeration to promote active filtration of the bacteria (24 h post-infection, 24 hpi); (ii) post-infection: challenged juveniles were taken out the infection tanks for 8 h at RT to ensure bacterial internalization within the pallial cavity (32 hpi). After this, juveniles were transferred to fresh tanks containing 200 mL of FSW with aeration and checked each 12 h (44 hpi, 56 hpi, 68 hpi…). Seawater was renewed daily (or whenever turbid), and clams were maintained without food throughout the second challenge period.

### 2.3 Survival rates and sample collection for DNA extractions

Survival rates were calculated and expressed as the percentage of surviving individuals (%) during both priming and second challenge steps (Figure 1). During the priming phase, survival was recorded at 0, 12, 24, 36, 48, 60, 72, 96, 120, 144, 168, and 192 hours post-priming (hpp) in both primed and non-primed groups (Figure 1). After the second challenge, mortality was recorded at 24, 32, 44, 56, 68, and 80 hpi. Because high survival was observed in primed clams following the second challenge, survival rate was also evaluated at 164 hpi (Figure 1). Clams were considered dead when their valves remained open, or moribund when siphons failed to retract upon stimulation. Dead and moribund juveniles were immediately removed from the tanks. Survival rates were plotted using Kaplan–Meier survival curves generated in Graphpad software. Statistical comparisons among curves were performed using Log-rank (Mantel-Cox) test for trend in multiple comparison analysis with the significance level set at p < 0.05.

**Figure 1.**
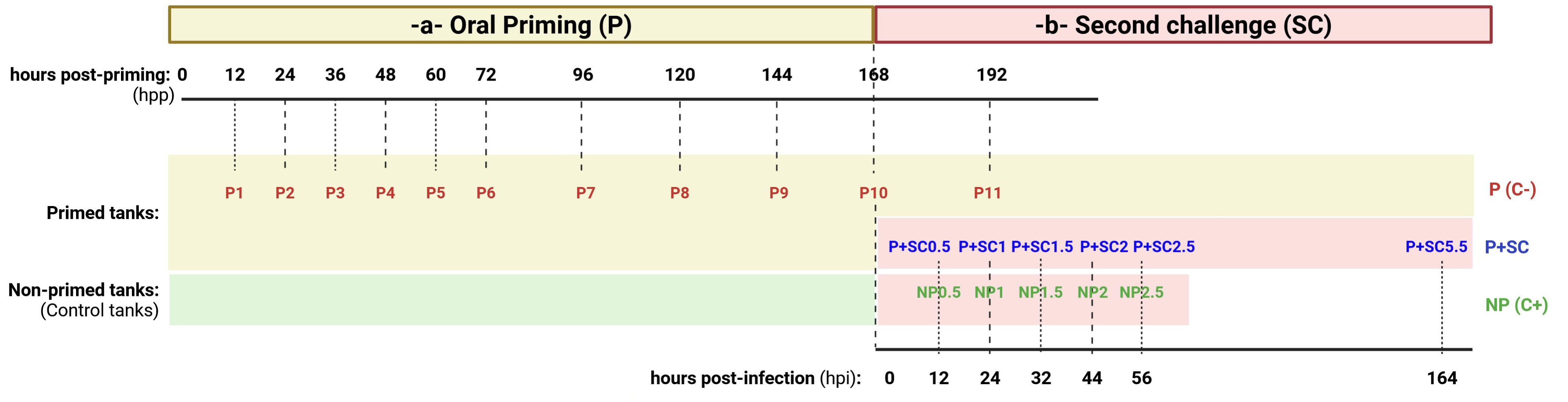
Overview of the experimental design used to evaluate clams survival (%) and sample collection for DNA extraction (pathogen clearance and full-length 16S metabarcoding). Sampling points are indicated as hours post-priming (hpp) or hours post-infection (hpi). Experimental conditions are represented by colors: (a) Red, animals primed during the first non-lethal priming stage (P0–P11); samples P10 and P11 were included for comparative analyses with those challenged during the second exposure and served as negative controls (P C–); (b) Second challenge, animals exposed to the bacterial pathogen during the second infection stage—blue, primed animals (P+SC; P+SC0.5– P+SC 5.5); and green, non-primed animals (NP C+; NP0.5–NP2.5).

Samples for total DNA extraction were collected at the timepoints indicated above (Figure 1). During the priming exposure, eleven samples (P1–P11) were collected from 0 to 192 hpp (Figure 1). Each sample consisted of a pool of three juveniles collected from each tank. Sample P11 (equivalent to 24 hpi), obtained from primed clams that were not infected during the second challenge, was used as a negative control for comparison with infected samples (Figure 1). After the second challenge, five samples were collected at 12, 24, 32, 44, and 56 hpi from primed clams, including both previously primed (P+SC0.5–P+SC2.5) and non-primed individuals (C+; NP0.5–NP2.5). Additionally a final sample for primed clams was collected at 164 hpi (P+SC5.5). For sampling, clam shells were individually removed using sterile scalpel blades, and the whole bodies were lyophilized and subsequently homogenized by bead beating using a Mixer Mill MM400 (Retsch). The resulting clam powder was stored at –80 °C until further use. Total DNA was extracted from approximately 1.5 mg of each sample using the DNeasy Blood and Tissue Kit (Qiagen) according to the manufacturer’s instructions. The concentration and purity of the extracted DNA were assessed using a NanoDrop One spectrophotometer (Thermo Scientific), a Qubit fluorometer (Thermo Scientific), and by capillary electrophoresis with an Agilent BioAnalyzer 2100 (Agilent Technologies).

### 2.4 Quantification of *V. europaeus* in challenged juveniles

Total DNA was used to evaluate pathogen clearance by the host during both the priming and second challenge exposures (Figure 1). The concentration of *V. europaeus* was quantified using a *ktrA*-based real time quantitative PCR (qPCR) TaqMan assay described by Rey-Varela et al. (2025). Primers, TaqMan probe and standard DNA nucleotide sequences are provided on Table S1. Each sample was analyzed in quadruplicate (two undiluted and two 10-fold diluted). Briefly, qPCR reactions were carried out in 20 µL containing 2 µL of template DNA, 10 µL Luna Universal Probe qPCR Master Mix (New England Biolabs), 0.4 µL Antarctic Thermolabile UDG (New England Biolabs), 0.4 µL probe (10 µM stock; final concentration 0.2 µM), 0.8 µL of each primer (10 µM stock; final concentration 0.4 µM), and 6.4 µL molecular-grade water. Reactions were run in 96-well plates sealed with heat-bonding film on a C1000 Touch thermocycler coupled to a CFX96 Touch Real Time PCR Detection System (Bio-Rad). Thermal cycling conditions consisted of an initial step at 25 °C for 10 min, followed by one cycle at 95 °C for 1 min, and 40 amplification cycles 95 °C for 15 s, 55°C for 15 s, and 68°C for 10 s. A standard curve was generated to determine the assay sensitivity, limit of detection and quantification, and amplification efficiency, using serial 10-fold dilutions of standard DNA (Table S1). qPCR results were expressed as *V. europaeus* genome copies per milligram of whole-body clam tissue (cop mg^−1^).

### 2.5 Library preparation and Full length 16S sequencing

PCR amplification, library preparation and sequencing were performed by Novogene (Cambridge, UK). For each sample, three independent amplification rounds were carried out using total DNA to amplify the full-length 16S rRNA gene with barcoded forward (27F: 5’-AGAGTTTGATCCTGGCTCAG-3’) and reverse primers (1510R: 5’-GGTTACCTTGTTACGACTT-3’), following the protocol described in Preparing Kinnex™ libraries from 16 rRNA amplicons (PacBio). The Kinnex 16S rRNA kit was used to increase throughput on PacBio long-read sequencers by applying a concatenation method that joins genomic DNA molecules (MAS-Seq) into longer fragments (Al’Khafaji et al., 2024).

Thermal cycling conditions consisted of an initial denaturation at 95 °C for 3 min, followed by 20 amplification cycles 98 °C for 20 s, 57°C for 30 s, and 72°C for 75 s, with a final extension at 72°C for 5 min, using 2X KAPA HiFi HotStart ReadyMix polymerase (Roche). The expected amplicon size (∼1,500 bp) was verified by electrophoresis on 1% (w/v) agarose gel (NZYtech) stained with GelRed Nucleic Acid Gel Stain (Biotium). Three replicates were pooled equimolarly and cleaned using SMRTbell Cleanup beads (PacBio) according to the manufacturer’s protocol. Cleaned PCR products were quantified using Qubit fluorometer (Invitrogen) with the 1x dsDNA HS Assay kit (Invitrogen). Subsequently, 16S amplicons were processed to prepare the Kinnex™ libraries following the PacBio protocol. This step adds Kinnex adapters to the ends of barcoded 16S full-length amplicons, enabling concatenation of PCR products to approximately19 kb. Library concentration was verified with Qubit fluorometer, and quantified libraries pooled and sequenced on a PacBio Sequel II/IIe system (Pacific Biosciences), according to effective library concentration and data amount required.

### 2.6 Microbiome taxonomical annotation

The PacBio BAM file was split according to barcode and filtered to get clean data. Briefly, PacBio offline data was exported to a BAM format file. Lima software was used to distinguish the data of each sample based on barcode sequences and save all sample sequences in BAM format. Then use CCS (SMRT Link v7.0) to correct the sequence, with a correction parameter of CCS=3 and a minimum accuracy of 0.99. Sequences with lengths less than 1340 and longest sequences with lengths greater than 1640 were removed, and stored in fastq and fasta; Subsequently, SSR filtration was performed and the primers were removed using cutadapt to filter out sequences containing consecutive identical base numbers>8. The Reads obtained after the above processing are the final valid data (Clean Reads) and shown in Table S2.

The obtained demultiplexed PacBio fastq sequence files were imported into QIIME2 (vQIIME2-202006). Denoise was performed with DADA2 or deblur module in the QIIME2 software (vQIIME2-202006) to obtain initial ASVs (Amplicon Sequence Variants) (default: DADA2), and then ASVs with abundance less than 5 were filtered out (Li et al., 2020). The ASVs were compared with the Silva 138 database(Quast et al., 2012). Rarefied ASV abundance was used for all downstream analyses (rarefied to 8441 reads). ASVs with fewer than 20 counts across all samples or present in only one condition were filtered out. The most 150 abundant ASVs were taxonomically assigned using EZBioCloud 16S-based ID (Chalita et al., 2024). ASVs assigned to the same taxon were subsequently collapsed into a single representative ASV, as detailed in Table S3.

Shannon index to obtain the alpha diversity (within samples) was calculated for all the samples with the R library vegan (v2.6-10) with the diversity command with index=“shannon” using default values. Bray-Curtis dissimilarity was calculated to obtain the beta diversity (between samples) with the R library vegan (v2.6-10) with the vegdist command with distance = “bray” using default values. Nonmetric Multidimensional Scaling (NMDS) was calculated from Bray-Curtis dissimilarity using the R library vegan (v2.6-10) with the command metaMDS, indicating distance=”bray” and k=2.

### 2.7 Normalization and clustering

Pseudocounts were obtained adding 1 to all the rarefied ASVs counts. Rarefied ASV pseudocount were normalized using the centered log-ratio (CLR) transformation with the R library compositions (v2.0-8) with the command clr using default values. To calculate the clusters, the distance between clr-normalized values using R library compositions (v2.0-8) with the command dist with method=”euclidean” using default values. The clustering was calculated using the command hclust with method = “ward.D2” using the default values. The hierarchical clustering was divided using the command cutree with k=6.

### 2.8 Ethical statement

Experimental work on invertebrates is not subjected to any ethical approval. However, all the challenges in this study were performed in accordance with good ethical and scientific practice.

## 3 Results

### 3.1 Oral priming effectively protects Manila clam against the bacterial pathogen *V. europaeus*

No mortalities were observed during the priming phase. However, a significant difference in survival (p<0.0001) was detected between primed (P+SC) and non-primed (NP C+) clams following the second challenge with *V. europaeus* at lethal dose (Figure 2). NP clams (C+) exhibited 100% mortality within 56 hours post-infection (hpi), whereas P+SC clams showed only minor mortalities starting at 68 hpi and maintained a stable survival rate of approximately 87% thereafter (Figure 2). Among primed clams, significant differences (p= 0.0117) were observed between those infected in the second challenge (P+SC) and non-primed (NP C–), which maintained 100% survival throughout the experiment (Figure 2). Overall, these results provide the first phenotypic evidence of oral immune priming in Manila clams, demonstrating effective protection against bacterial infection by *V. europaeus*.

**Figure 2.**
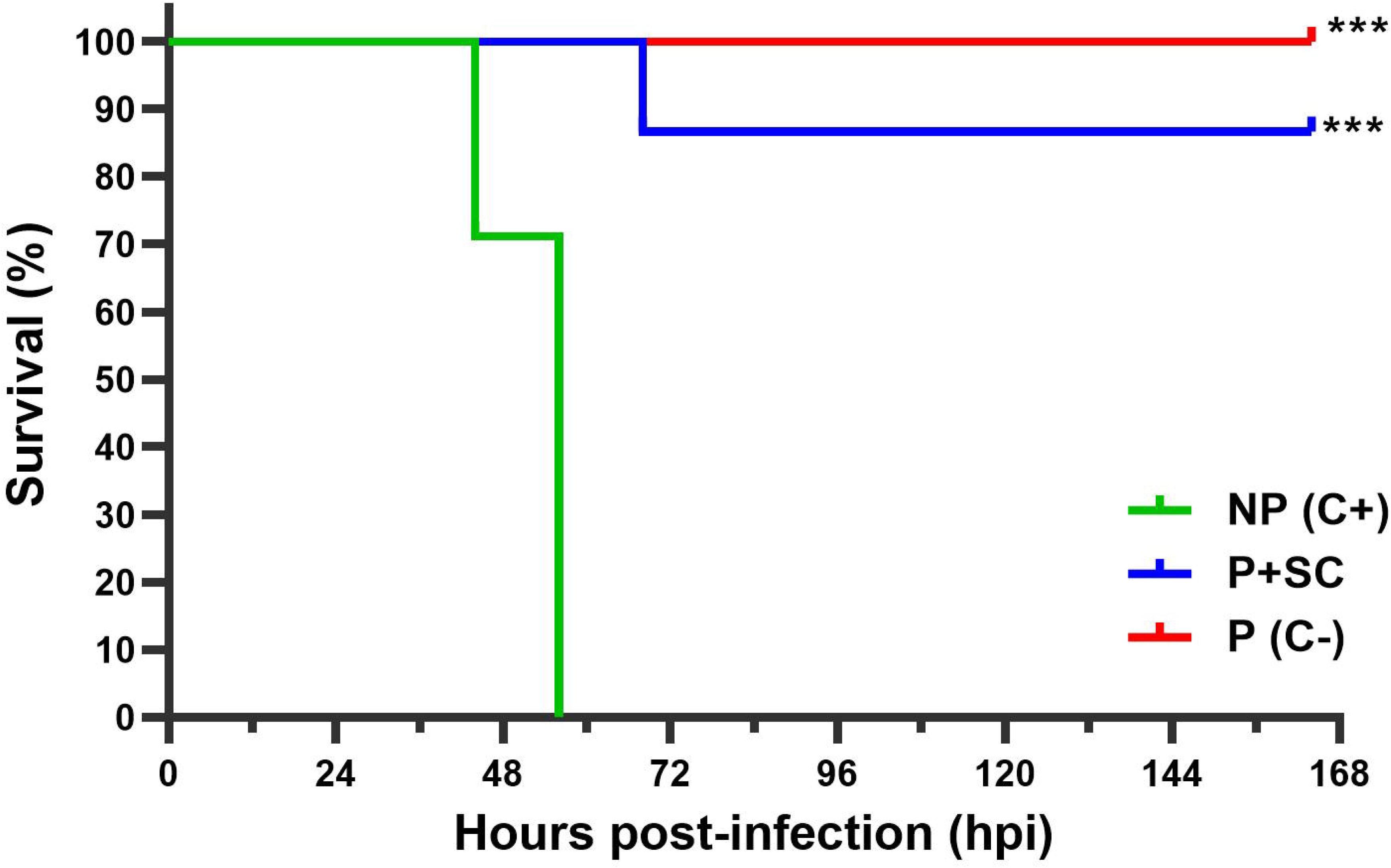
Kaplan Meier survival curves (%) of Manila clam juveniles following second challenge. Primed clams (P+SC, blue curve) and non-primed clams (NP C+, green curve) were infected with *V. europaeus* at a lethal dose. Primed clams that non-challenged with the pathogen served as negative controls (P C-, red curve). Asterisks indicate significant differences with the non-primed group (NP - C+) (p<0.001).

### 3.2 Primed clams reduce the bacterial pathogen to similar concentrations after both priming and second challenge

During the priming phase (Figure 3A), the concentration of *V. europaeus* decreased by two orders of magnitude—from 3.21 × 10^4^ to 4.15 × 10^2^ copies mg^−1^—within the first 48 hours post-priming (hpp) and remained stable throughout the experiment, including in primed clams that were not challenged after 192 hpp (equivalent to 24 hpi; P C–).

**Figure 3.**
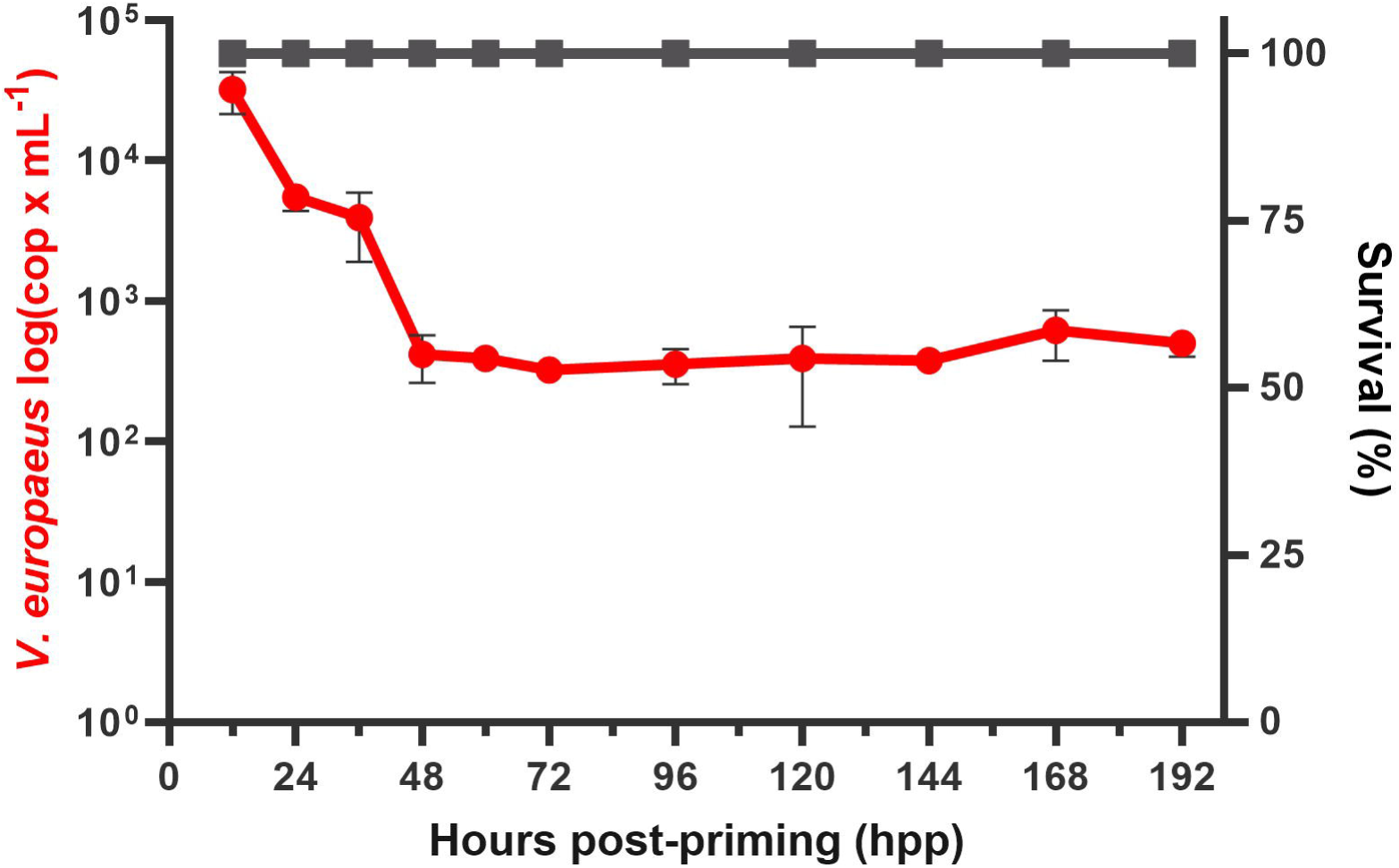
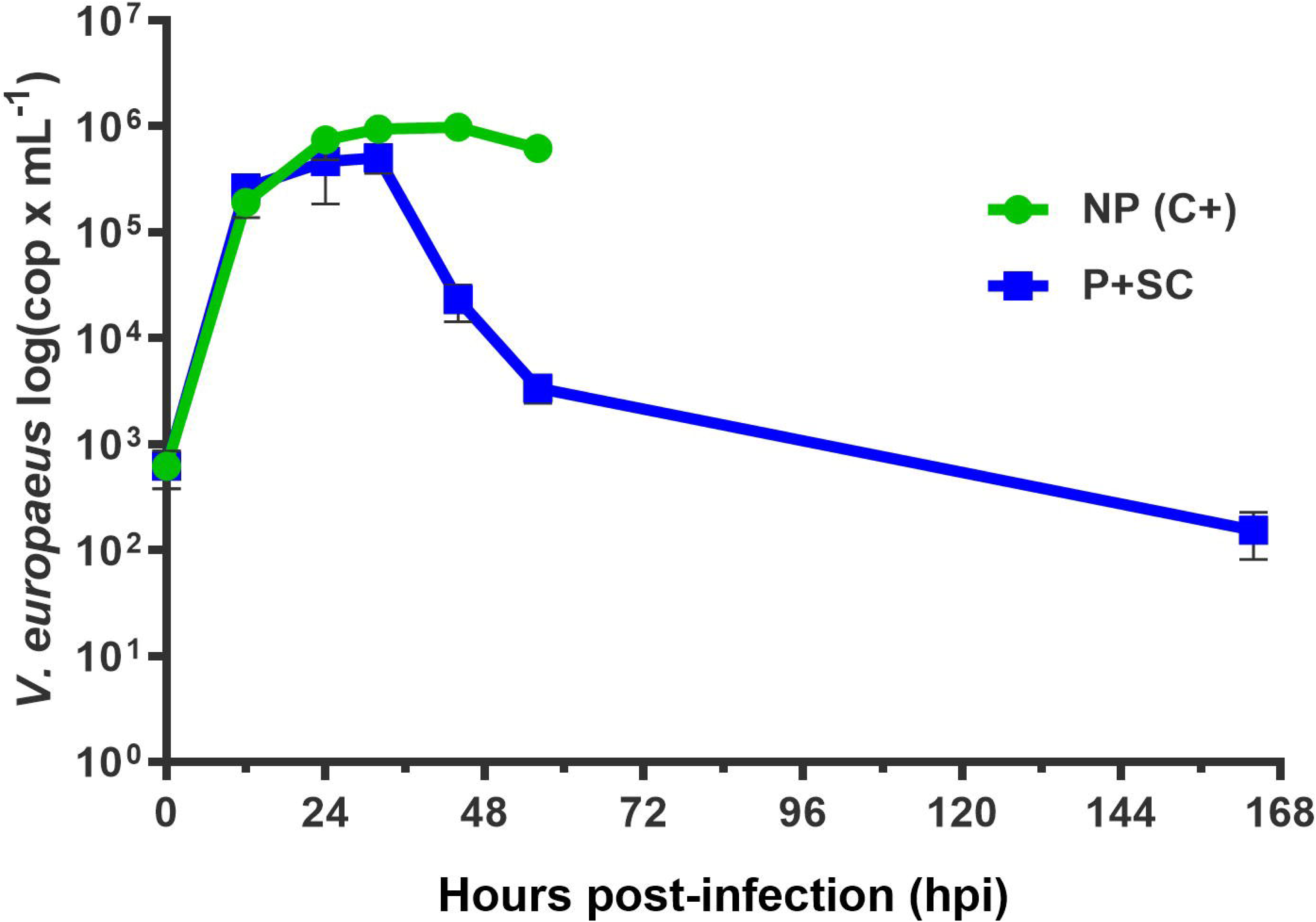
Pathogen clearance during priming (A) and second challenge (B) phases. A, quantification of *Vibrio europaeus* expressed as genome copies per milligram of clam tissue determined by qPCR. The red line represents pathogen concentrations during oral priming. The grey line represents survival during the priming phase, in which no mortalities were detected. B, pathogen concentration following the second challenge in primed clams (P+SC, blue line) and non-primed clams (NP, C+; green line).

In P+SC clams, pathogen concentrations increased during the first 32 hours, reaching a peak of 5.07 × 10^5^ copies mg^−1^ (Figure 3B). However, P+SC clams reduced bacterial concentrations by approximately two orders of magnitude from that time, reaching 3.37 × 10^3^ copies mg^−1^ at 56 hpi. Remarkably, concentrations further decreased to 1.54 × 10^2^ copies mg^−1^ at 164 hpi, levels comparable to those observed during the priming phase.

In contrast, NP (C+) clams were unable to reduce bacterial concentrations below 10⁵ copies mg^−1^ after 32 hpi, which coincided with the high mortalities observed in this group and subsequent death of the individuals (Figure 2 and 3B). These results highlight (i) the capacity of primed clams to control and reduce *V. europaeus* concentration below the threshold (∼10^5^ copies mg^−1^ after 32 hpi) associated with high mortality in non-primed animals, and (ii) the persistence of the pathogen at low concentrations in primed clams without apparent harm to the host.

### 3.3 Alpha diversity reveals that priming reduces the microbiota diversity

The microbiota composition of all experimental groups was analyzed using full-length 16S rRNA gene sequencing. Amplicon sequence variants (ASVs) belonging to the same species were collapsed using the Eztaxon database, resulting in a total of 492 ASVs identified across all experimental groups. Overall microbiota diversity was evaluated through alpha diversity analyses (Figure 4A). During the priming phase, alpha diversity in primed clams progressively declined over time, with a marked reduction between 120 and 192 hpp. The lowest diversity values were observed at 144 hpp (P9) and 192 hpp (P11).

**Figure 4.**
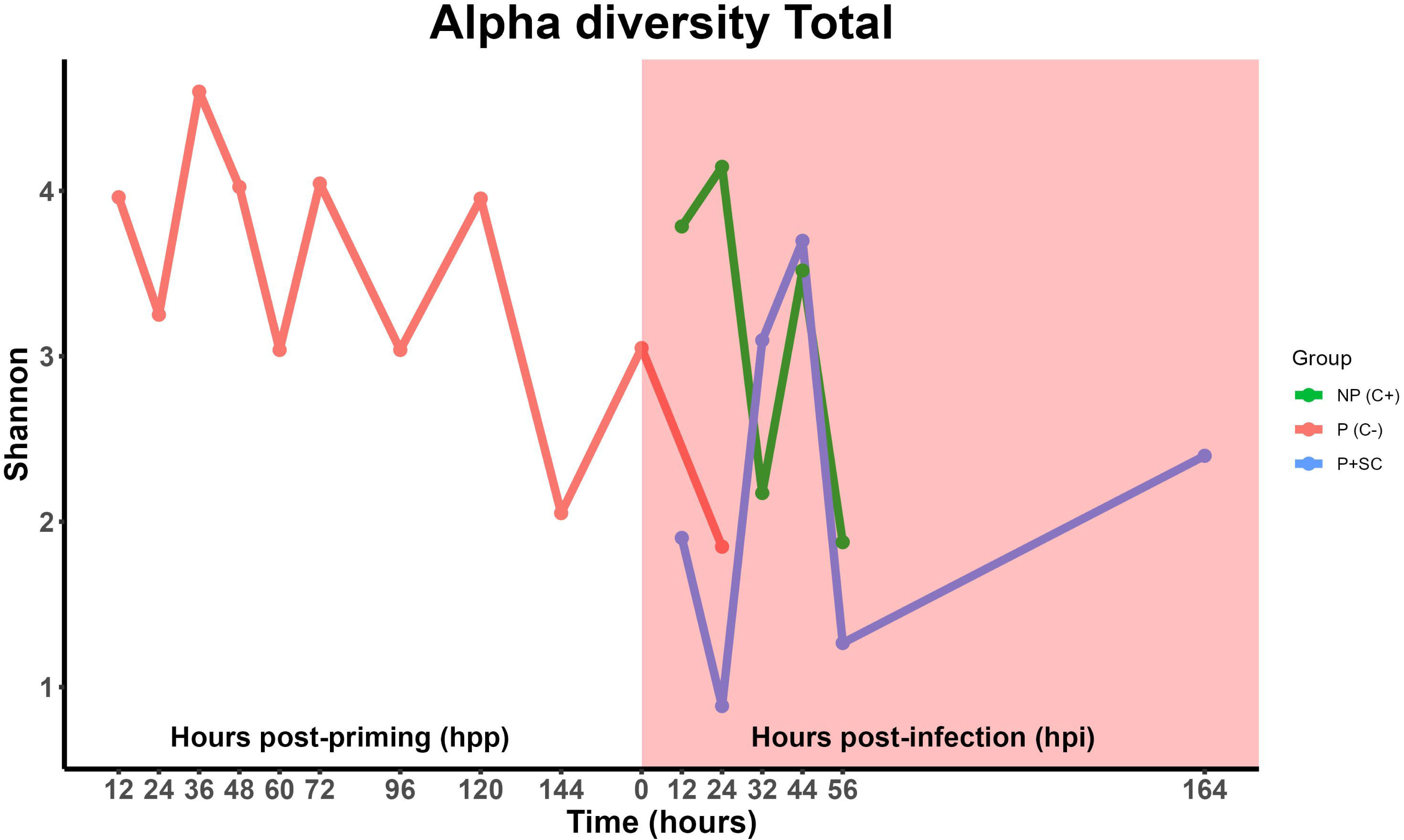
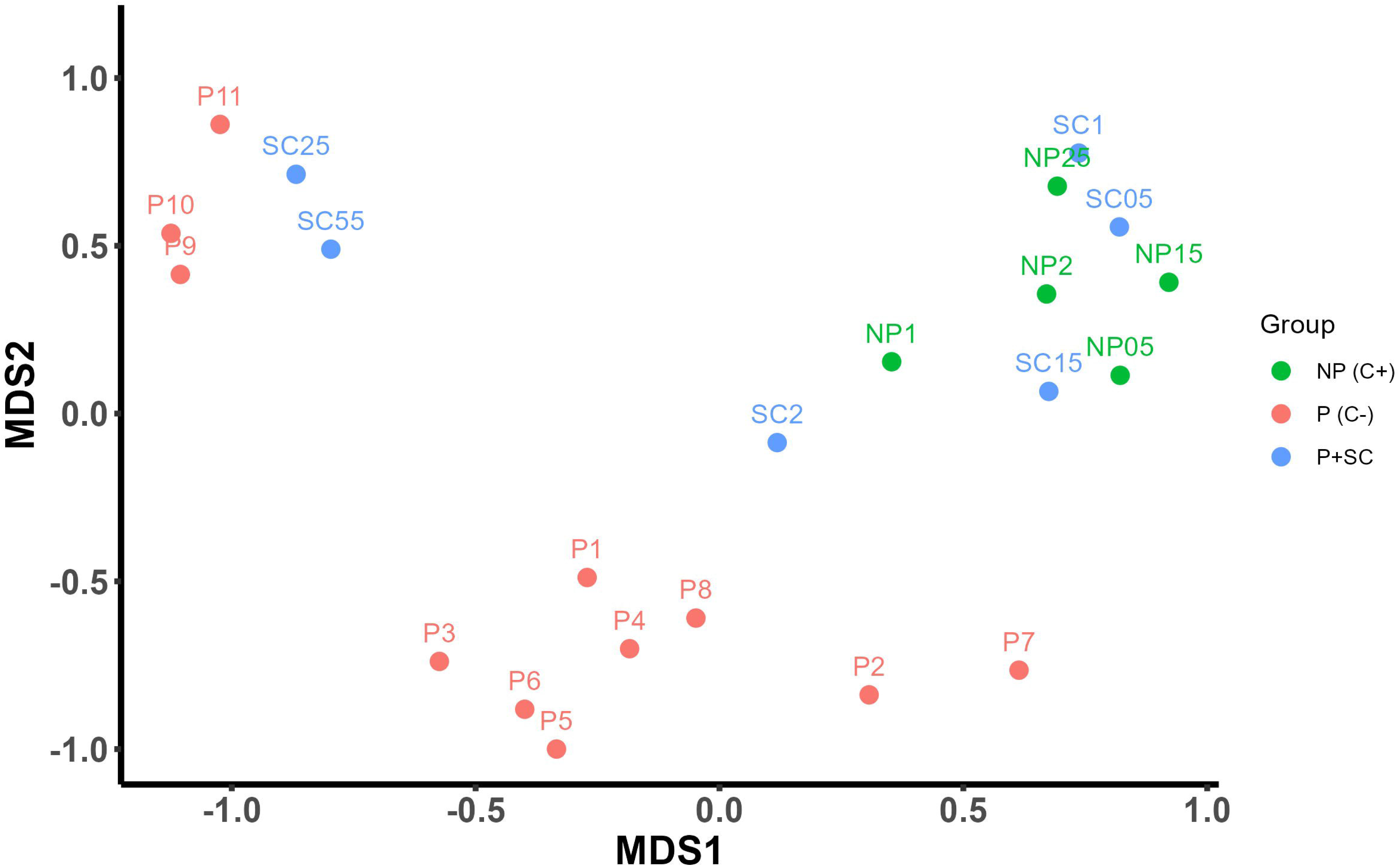
Alpha (A) and beta (B) diversity analyses of the Manila clam microbiota during the priming and second challenge phases. A, Alpha diversity (Shannon’s index) across experimental groups during the priming phase and the second challenge. Variations in diversity are shown for primed clams (P, red line), primed and subsequently challenged clams (P+SC, blue line), and non-primed clams (NP, green line) (pink shaded area indicates the time points corresponding to the second challenge). B, Beta diversity visualized by multidimensional scaling (MDS) based on Bray–Curtis dissimilarities. Samples collected during the priming phase (P1–P11) and the second challenge (P+SC and NP) are shown.

Following the second challenge, *V. europaeus* infection triggered an antagonistic diversity dynamic between P+SC and NP clams during the first 32 hpi (Figure 4A). Specifically, NP clams exhibited a sharp increase in alpha diversity, reaching the highest peak at 24 hpi, whereas P+SC clams showed a pronounced decrease, attaining the lowest diversity at the same time point (Figure 4A). From 24 to 32 hpi, these trends diverged further: diversity progressively decreased in NP clams but gradually increased in P+SC clams. This antagonistic response suggests a direct effect of the initial priming exposure on the modulation of bacterial community diversity and host resistance during the first 32 hpi (Figure 4A). After 44 hpi, diversity dynamics converged between both groups and progressively decreased until 56 hpi. In NP clams, this reduction coincided with increased mortality and higher pathogen loads observed by experimental challenges and qPCR respectively, suggesting dominance by *V. europaeus* (Figures 2, 3B, and 4A). In contrast, the decline in diversity observed in P+SC clams paired with a reduction in pathogen concentration (Figure 3B), supporting the proliferation of specific microbial communities associated with the immune priming response. Interestingly, by the end of the second challenge (164 hpi), alpha diversity in P+SC clams recovered to levels comparable to those observed at the end of the priming phase (P9–P11), indicating partial restoration of the microbiota structure following infection.

### 3.4 Beta diversity demonstrates the establishment of a specific community composition associated with immune priming

To assess compositional differences, Bray–Curtis dissimilarities were calculated and visualized via multidimensional scaling (MDS) (Figure 4B). During the priming phase, samples from 12–120 hpp (P1–P8) clustered closely, but from 144 hpp (P9) onward, the microbiota composition shifted and then stabilized, clustering samples P9, P10 and P11 together (Figure 4B).

Beta diversity analysis revealed clear compositional distinctions between priming and secondary challenge stages. During the initial 32 hpi, both P+SC and NP samples (P+SC0.5–1.5; NP0.5–1.5) clustered together, coinciding with a sharp increase in pathogen abundance (∼10^5^ copies mg^−1^; Figure 3B). NP samples collected at 44 and 56 hpi (NP2 and NP2.5) also grouped within this cluster, consistent with peak mortality events (Figure 2). Conversely, from 44 hpi onward, only P+SC clams (PSC2–5.5) gradually returned to a community structure comparable to the end of the priming phase, clustering again with samples P9–P11, coinciding with a two-log reduction in pathogen load (Figure 3B and 4B).

Together, these findings indicate that oral priming induces a distinct and stable microbiota composition in Manila clams. This primed community exhibits enhanced resilience and the capacity to recover following subsequent *V. europaeus* infection, suggesting a microbiota-mediated contribution to host resistance.

### 3.5 Immune priming promotes a specific and stable microbiota

Clustering analysis was performed to identify the bacterial taxa associated with primed clams, which grouped together in beta diversity plot at the end of both the priming phase (P9–P11) and the second challenge (SL2.5–SL5.5). The microbiota of these samples was compared with that of NP clams (NP0.5–NP2.5). After CLR-normalization and hierarchical clustering, elbow method was used to determine that, the optimal number of clusters is six (Supplementary Figure S1).

Clusters 1 and 5 (Figure 5) best explained the overall beta and alpha diversity patterns observed. In Cluster 5, primed clams at the end of the second challenge (PSC2.5 and PSC5.5) harbored a specific microbiota dominated by Alphaproteobacteria, including *Sphingobium limneticum* (ASVB12) and *Brevundimonas* (*B. huaxiensis/B. vesicularis/B. nasdae/B. fontaquae/B. intermedia*; ASVB13), as well as Gammaproteobacteria belonging to the family *Moraxellaceae*, such as *Acinetobacter* (*A. lwoffii/Prolinoborus fasciculus/A. pecorum*; ASVB7) and *Psychrobacter alimentarius* (ASV37). All these taxa were also detected in clams at the end of the priming phase (P9–P10) and especially in the sample corresponding to the onset of the second challenge (P10, 0 hpi) (Figure 5). Particularly *Acinetobacter* and *Brevundimonas* were also identified in the negative control corresponding to 12 hpi (P11; Figure 5). Their progressive proliferation in PSC2.5–PSC5.5 supports the hypothesis that these groups contribute to the microbiota stability induced by priming since members of Cluster 5 were absent or progressively declined in NP clams.

**Figure 5.**
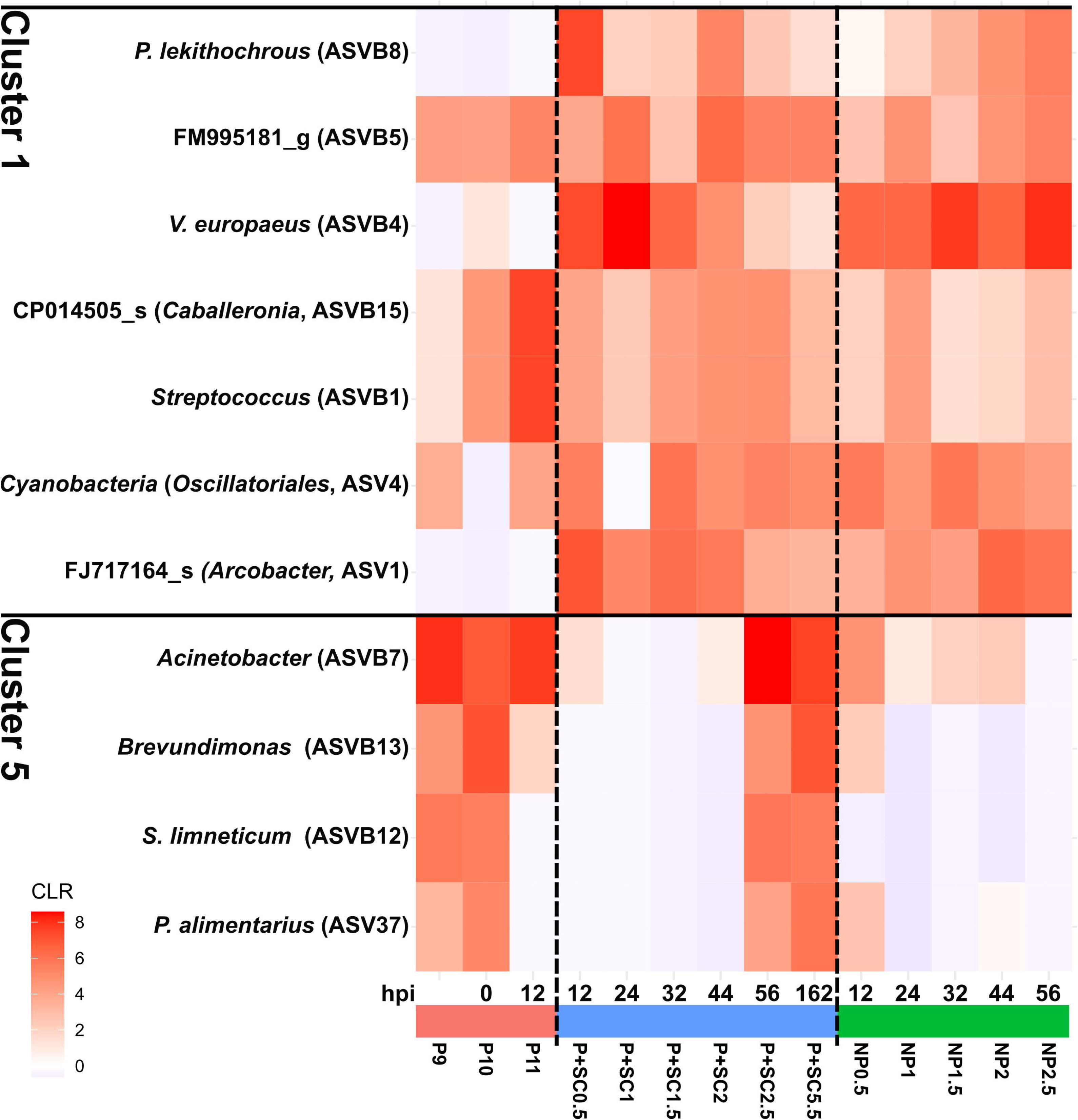
Clustering analysis of bacterial taxa associated with the Manila clam microbiota during the priming and second challenge phases. Rarefied amplicon sequence variant (ASV) counts were normalized using the centered log-ratio (CLR) transformation. Clustering analysis based on CLR-normalized ASV abundances identified six major clusters of bacterial taxa across the experimental groups. The heatmap shows the bacterial composition corresponding to Cluster 1 and Cluster 5, which are the clusters that best explain the beta diversity distribution (clusters 2, 3, 4 and 6 are shown in the supplementary material).

In relation to Cluster 1, several bacterial taxa were absent in P clams after priming but appeared during the second challenge, likely in association with infection. These taxa—excluding the bacterial pathogen *V. europaeus* (ASVB4)—increased in NP clams but decreased in primed clams and included members of the family *Arcobacteraceae*, such as *Poseidonibacter lekithochrous* (ASVB8) and an unidentified species (FJ717164_s) belonging to the genus *Arcobacter* (ASV1) (Figure 5).

Other taxa belonging to Cluster 1 were shared among PSC, NP, and P clams throughout the experiment, suggesting that they constitute part of the core or resident microbiota (Figure 5). In Cluster 1, these included *Streptococcus* (*S. thermophilus/S. salivarius/S. vestibularis*; ASVB1) and unidentified bacteria such as FM995181_g (ASVB5), CP014505_s belonging to genus *Caballeronia* (ASVB15), and an unknown cyanobacterium assigned to the order Oscillatoriales (ASV4) (Figure 5). This was also observed in some taxa assigned to Cluster 4, it included to *Cutibacterium acnes* (ASVB9), an unknown species belonging to genus *Oceaniferula* (GQ262724s; ASV32), *Cellulophaga* (ASV50) and *Phaeobacter* (ASV48) (Supplementary Figure S2).

Clusters 3 and 6 (Supplementary Figure S3 and S4) showed the taxa involved in the high alpha diversity observed in NP clams at 12 hpi (Cluster 3) and 24 hpi (Cluster 6), demonstrating that both clusters are clearly associated to infection and suggesting that could not be associated to primed clams due to any unknown mechanisms related with immune priming to avoid the proliferation of those taxa. Finally, cluster 2 showed taxa punctually detected in the samples analyzed (Supplementary Figure S5). Overall, these results demonstrate the existence of a specific and stable microbiota associated with the immune priming response.

## 4 Discussion

This study provides the first phenotypic evidence of successful oral immune priming in the Manila clam (*Ruditapes philippinarum*), demonstrating a robust protection against the bacterial pathogen *V. europaeus*. From the perspective of the priming stimulus, many studies have relied on inactivated pathogens, it means the use of dead cells to induce the immune priming response in the hosts (Y. Li et al., 2017; Liu et al., 2016; J. Wang et al., 2013; Yue et al., 2013; T. Zhang et al., 2014). However, in natural environments, priming responses are expected to occur in response to live pathogens. Thus, our experimental approach relies on the use of live cells of the pathogen at a sublethal dose to induce the priming effect, followed by a second challenge at lethal concentrations to evaluate protective capacity. This experimental approach allows the possibility to study pathogen dynamics after priming and after second challenge through its quantification by qPCR. Clams during the priming step effectively reduced the *V. europaeus* concentration by two orders of magnitude within 48 hours post-priming (hpp) and maintained stable pathogen levels similar to those observed in primed clams after the second challenge (approximately 10^2^ cop mg^−1^). This rapid decrease reduced pathogen load below the critical threshold associated with mortality (∼10⁵ copies mg⁻¹), resulting in the survival of primed individuals. In contrast, non-primed clams also reached pathogen concentrations of ∼10⁵ copies mg⁻¹, which were not reduced and coincided with the massive mortalities (100%) observed in this group. These findings confirm the persistence of innate immune memory in a key aquaculture bivalve such as the Manila clam, extending previous evidence reported (S. Li et al., 2025).

Immune priming in bivalves has mostly been induced by injection, and particularly in Manila clam (Lafont et al., 2017, 2020; S. Li et al., 2025; Rey-Campos et al., 2019; J. Wang et al., 2013; T. Zhang et al., 2014). In contrast, this study demonstrates that oral administration—both for priming and challenge—can successfully induce immune protection. The rationale lies in that bivalves are filter feeders, and thus, the oral route is their natural route of infection (Dubert et al., 2017). Moreover, injection-based approaches induced physiological and immunological biases due to tissue trauma, which triggers itself an immune activation, for instance, the wounding intrinsic to injection induces itself a strong immune response (Behrens et al., 2014; Johnston & Rolff, 2013). For Manila clam, the oral route yielded higher survival rates (87%) than those obtained in previous studies by injection (82%) (S. Li et al., 2025), and notably, in oral priming model no mortality occurred during the priming phase. In contrast, the injection-based study reported 69% mortality during priming (S. Li et al., 2025), likely due to wounding stress and unnatural exposure conditions such as the high bacterial concentration used at this step (5 × 10^7^ CFU ml^−1^). The oral route thus provides a more physiologically relevant and less invasive strategy to induce immune priming in bivalves and minimizing host stress.

From an applied perspective, oral immunization also offers clear practical advantages: it enables the possibility to immunize a high number of animals without the need for handling individual animals, making it particularly suitable for large-scale aquaculture systems. Despite decades of intensive mollusk production, no effective, eco-friendly prophylactic or therapeutic strategies have yet been established to fight bacterial pathogens in shellfish aquaculture (Dubert et al., 2017a; Smolowitz, 2024). Immune priming therefore emerges as a promising approach to address this gap. The survival rates achieved in this study are comparable to —or higher than— those obtained in other immune priming studies in different mollusk hosts towards different *Vibrio* species (Montagnani et al., 2024). For example, effective protection has been achieved by injection-based priming with *V. anguillarum* and *V. parahaemolyticus* (J. Wang et al., 2013; X. Zhang et al., 2022), as well as by immersion with *V. harveyi* in abalone (Dubief et al., 2017) and formalin-killed *V. alginolyticus* in *Magallana gigas* (H. Wang et al., 2024). Interestingly, in the latter case, only inactivated bacteria conferred protection, while live sublethal priming was ineffective—suggesting that priming mechanisms are highly context-dependent and may vary across host–pathogen systems (Lanz-Mendoza & Contreras-Garduño, 2022; Montagnani et al., 2024).

This study demonstrates for the first time in *Ruditapes philippinarum* that immune priming is accompanied by a profound shift of host’s microbiota both in bacterial diversity and community composition. During the priming phase, the microbiota displayed an initial increase in alpha diversity followed by a gradual, progressive reduction, with the lowest values recorded between 144 and 192 hpp. This narrowing of the microbial community suggests a selective process favoring a core, resilient bacterial community that emerges as a consequence of the priming stimulus. The structural bacterial divergence between treatments became particularly evident following the second challenge. Non-primed clams, which succumbed to infection, exhibited a sharp rise in alpha diversity—peaking at 24 hpi—a pattern typically associated with dysbiosis, reflecting loss of microbial homeostasis due to host immune failure and the opportunistic proliferation of multiple taxa. In contrast, protected clams showed a marked decrease in diversity at the same time point. This supports the idea that the priming-associated microbiota may function as a stabilizing bottleneck, counteracting the early invasive changes triggered by infection. Furthermore, Beta diversity analyses further confirmed the establishment of a distinct and stable community composition associated with the primed state. Protected clams progressively returned to the microbial structure established at the end of the priming phase (P9–P11), a compositional recovery that coincided with the two-log reduction in pathogen burden. The persistence and re-emergence of post-priming ASVs—particularly those in Cluster 5—after the acute stage of the second challenge reinforce the link between microbiota composition and priming-induced resilience.

Cluster 5 included taxa such as *Acinetobacter* and *Psychrobacter alimentarius*, which may contribute positively to the immune priming response. Although *Acinetobacter* species are often recognized for their pathogenic potential, some isolates show antimicrobial activity against *Vibrio* spp. in Seriola aquaculture (Ramírez et al., 2020). Similarly, *Psychrobacter* sp. B6 has displayed probiotic activity against *Aeromonas hydrophila* (Ramírez et al., 2020). Our findings support the idea of a reciprocal interaction: the priming stimulus shapes the microbial community, and in turn, this microbiota specifically associated with immune priming contributes to the host protection. Interactions between microbiota and immune priming have been mostly studied in arthropods, but are largely overlooked in marine mollusks (Montagnani et al., 2024). In insects, microbiota plays a pivotal role in activating immune priming. For instance, removal of the gut microbiota in mosquitoes (*Anopheles gambiae*) abolished immune priming protection against *Plasmodium falciparum* (Rodrigues et al., 2010). Similar microbiota-dependent immune priming responses have been observed in beetles, moths, honeybees, and cockroaches, where symbiotic microorganisms modulate host immunity by altering bacterial composition or abundance, inducing immune effectors (e.g., antimicrobial peptides), or regulating immune signaling pathways (e.g., Toll pathway) (Futo et al., 2016; Hernández-Martínez et al., 2010; Horak et al., 2020; Jang et al., 2024; Keshavarz et al., 2025; Korša et al., 2022; Lang et al., 2022; Turner et al., 2024). In marine invertebrates, however, the role of microbiota in immune priming remains largely unexplored. The only example reported so far is in the Pacific oyster (*M. gigas*), where exposure to non-infectious environmental microbiota during early larval stages (larvae) induced a systemic immune response that conferred disease protection (Fallet et al., 2022). The protection observed in our study is therefore likely linked to a profound restructuring of the host microbial community. By focusing specifically on the microbiota associated with immune priming response, we demonstrate that oral immune priming establishes a specific and resilient microbiota composition in the Manila clam. While previous molluscan studies have concentrated primarily on host immune responses (Montagnani et al., 2024), our findings suggest that the microbiota plays an active and central role in the priming process in molluscs.

Because Cluster 1 contains *V. europaeus*, its dynamic suggests a mechanism in which the host’s immune system, reinforced by priming, decrease the proliferation of the pathogen and possibly other co-infecting species such as members of the family *Arcobacteraceae*. However, this bacterial dynamic was opposite in non-primed animals. Members of the genus *Arcobacter* have been identified together with *Vibrio* as pathobionts that play a major role in the development of Pacific Oyster Mortality Syndrome (POMS), a significant threat to the aquaculture industry (Clerissi et al., 2023). POMS involves profound changes in the oyster microbiota in response to infection with the OsHV-1 µVar virus. Infection by OsHV-1 µVar induces an immunocompromised state that triggers microbiota dysbiosis and subsequent bacteremia, ultimately leading to oyster death. Fatal dysbiosis is accompanied by invasion of connective tissues by opportunistic bacteria—particularly *Vibrio* and *Arcobacter*—which are consistently associated with mortality events (Clerissi et al., 2023). In our case, these findings highlight that *V. europaeus* proliferation in disease clams is accompanied by the proliferation of other bacterial taxa, particularly *Arcobacter*. Although members of the genus *Arcobacter* are known to cause human gastroenteritis associated with the consumption of raw bivalves, their role in bivalve pathogenesis remains poorly understood (Mottola et al., 2016; Ramees et al., 2017). For example, *Poseidonibacter lekithochrous*, a member of the *Arcobacteraceae* associated to non-primed clams here, was previously reported by our group as a species not associated with mortality events in shellfish hatcheries (Diéguez et al., 2017). Other genera present in Cluster 1 have not previously been linked to mollusks—or to animals in general. However, further research will be needed to clarify their potential implications in disease progression during infection challenges.

From the exclusive perspective of the pathogen, it is important to remark the persistence of *V. europaeus* at low, non-harmful concentrations in primed clams suggests a state of controlled tolerance, rather than full pathogen elimination, a hallmark of a balanced host–pathogen interaction. Host responses to infections, such as resistance or tolerance, play a critical role in shaping virulence evolution. Recently, it has been published the first report showing the influence of insect specific immune priming on pathogen evolution using the red flour beetle (*Tribolium castaneum*) as model (Korša et al., 2025).Those authors revealed an increased activity in mobile genetic elements harbored by the pathogen (*Bacillus thuringiensis tenebrionis*), including prophages and plasmids, with variations in a virulence-related plasmid encoding the Cry toxin. This highlights that immune priming can promote diversity in pathogen traits, which may favor adaptation to variable environments. This underscores the need to consider pathogen evolution in response to immune priming when applying immune priming strategies in medicine, aquaculture, pest control, and insect mass production (Korša et al., 2025).

This study validates the oral delivery route as a viable method for inducing protection in Manila clam and suggests a crucial role for the microbiota as a central determinant of the immune priming response in bivalves. These findings open new avenues for the development of prebiotic and probiotic strategies where specific microbial communities could be manipulated to confer disease resistance—an urgently needed, as sustainable prophylactic strategy in aquaculture. Moreover, the results clearly demonstrate this intricate relationship, suggesting that the protective mechanism of immune priming cannot be fully explained by a simple bipartite interaction. Instead, we propose that the study of immune priming must shift towards a holistic approach from the study of the tripartite interactions between the host’s immune system, its microbiota, and the pathogen.

## 6 Author Contributions

JD, ALD and BKRJ conceived and designed the experiments. SR and CMS performed the experiments. JD, BKRJ, ALD and DRV analysed the data, wrote and revised the manuscript.

## 7 Acknowledgments

We are grateful for the financial support provided by the Spanish State Research Agency through the Grant CNS2022-136130 funded by MICIU/AEI/10.13039/501100011033 and by “European Union NextGenerationEU/PRTR”.

## 8 Data Availability Statement

The sequencing data were deposited at the NCBI Sequence Read Archive under BioProject PRJNA1358957. The ASV table with counts for each experimental condition is indicated in the supplementary file Supp_PrimingASVs.csv.

## References

Aleng, N. A., Sung, Y. Y., MacRae, T. H., & Wahid, M. E. A. (2015). Non-lethal heat shock of the asian green mussel, *Perna viridis*, promotes Hsp70 synthesis, induces thermotolerance and protects against *Vibrio* infection. PLOS ONE, 10(8), e0135603. 10.1371/JOURNAL.PONE.0135603

Al’Khafaji, A. M., Smith, J. T., Garimella, K. V., Babadi, M., Popic, V., Sade-Feldman, M., Gatzen, M., Sarkizova, S., Schwartz, M. A., Blaum, E. M., Day, A., Costello, M., Bowers, T., Gabriel, S., Banks, E., Philippakis, A. A., Boland, G. M., Blainey, P. C., & Hacohen, N. (2024). High-throughput RNA isoform sequencing using programmed cDNA concatenation. Nature biotechnology, 42(4), 582–586. 10.1038/s41587-023-01815-7

Behrens, S., Peuß, R., Milutinović, B., Eggert, H., Esser, D., Rosenstiel, P., Schulenburg, H., Bornberg-Bauer, E., & Kurtz, J. (2014). Infection routes matter in population-specific responses of the red flour beetle to the entomopathogen *Bacillus thuringiensis*. BMC Genomics, 15(1), 1–17. 10.1186/1471-2164-15-445/FIGURES/7

Chalita, M., Kim, Y. O., Park, S., Oh, H. S., Cho, J. H., Moon, J., Baek, N., Moon, C., Lee, K., Yang, J., Nam, G. G., Jung, Y., Na, S. I., Bailey, M. J., & Chun, J. (2024). EzBioCloud: a genome-driven database and platform for microbiome identification and discovery. International Journal of Systematic and Evolutionary Microbiology, 74(6). 10.1099/IJSEM.0.006421

Clerissi, C., Luo, X., Lucasson, A., Mortaza, S., de Lorgeril, J., Toulza, E., Petton, B., Escoubas, J. M., Dégremont, L., Gueguen, Y., Destoumieux-Garzόn, D., Jacq, A., & Mitta, G. (2023). A core of functional complementary bacteria infects oysters in Pacific Oyster Mortality Syndrome. Animal Microbiome, 5(1), 26-. 10.1186/S42523-023-00246-8/FIGURES/6

Cong, M., Song, L., Qiu, L., Li, C., Wang, B., Zhang, H., & Zhang, L. (2009). The expression of peptidoglycan recognition protein-S1 gene in the scallop *Chlamys farreri* was enhanced after a second challenge by *Listonella anguillarum*. Journal of Invertebrate Pathology, 100(2), 120–122. 10.1016/J.JIP.2008.10.004

Cong, M., Song, L., Wang, L., Zhao, J., Qiu, L., Li, L., & Zhang, H. (2008). The enhanced immune protection of Zhikong scallop *Chlamys farreri* on the secondary encounter with *Listonella anguillarum*. Comparative Biochemistry and Physiology Part B: Biochemistry and Molecular Biology, 151(2), 191–196. 10.1016/J.CBPB.2008.06.014

Diéguez, A. L., Balboa, S., Magnesen, T., & Romalde, J. L. (2017). *Arcobacter lekithochrous* sp. nov., isolated from a molluscan hatchery. International Journal of Systematic and Evolutionary Microbiology, 67(5), 1327–1332. 10.1099/IJSEM.0.001809/

Dubert, J., Nelson, D. R., Spinard, E. J., Kessner, L., Gomez-Chiarri, M., da Costa, F., Prado, S., & Barja, J. L. (2016). Following the infection process of vibriosis in Manila clam (*Ruditapes philippinarum*) larvae through GFP-tagged pathogenic *Vibrio* species. Journal of Invertebrate Pathology, 133, 27–33. 10.1016/j.jip.2015.11.008

Dubert, J., Barja, J. L., & Romalde, J. L. (2017a). New insights into pathogenic Vibrios affecting bivalves in hatcheries: present and future prospects. Frontiers in Microbiology, 8(MAY). 10.3389/FMICB.2017.00762

Dubert, J., Aranda-Burgos, J. A., Ojea, J., Barja, J. L., & Prado, S. (2017b). Mortality event involving larvae of the carpet shell clam Ruditapes decussatus in a hatchery: isolation of the pathogen *Vibrio tubiashii* subsp. *europaeus*. Journal of Fish Diseases, 40(9), 1185–1193. 10.1111/jfd.12593

Dubief, B., Nunes, F. L. D., Basuyaux, O., & Paillard, C. (2017). Immune priming and portal of entry effectors improve response to vibrio infection in a resistant population of the European abalone. Fish & Shellfish Immunology, 60, 255–264. 10.1016/J.FSI.2016.11.017

Fallet, M., Montagnani, C., Petton, B., Dantan, L., de Lorgeril, J., Comarmond, S., Chaparro, C., Toulza, E., Boitard, S., Escoubas, J. M., Vergnes, A., Le Grand, J., Bulla, I., Gueguen, Y., Vidal-Dupiol, J., Grunau, C., Mitta, G., & Cosseau, C. (2022a). Early life microbial exposures shape the *Crassostrea gigas* immune system for lifelong and intergenerational disease protection. Microbiome, 10(1), 1–21. 10.1186/S40168-022-01280-5/FIGURES/9

FAO. (2024). The State of World Fisheries and Aquaculture. Food and Agriculture Organization, 2024, 1–244.

Futo, M., Armitage, S. A. O., & Kurtz, J. (2016). Microbiota plays a role in oral immune priming in *Tribolium castaneum*. Frontiers in Microbiology, 6(JAN), 171102. 10.3389/FMICB.2015.01383/TEXT

Gerdol, M., & Venier, P. (2015). An updated molecular basis for mussel immunity. Fish & Shellfish Immunology, 46(1), 17–38. 10.1016/J.FSI.2015.02.013

Green, T. J., Helbig, K., Speck, P., & Raftos, D. A. (2016). Primed for success: Oyster parents treated with poly(I:C) produce offspring with enhanced protection against Ostreid herpesvirus type I infection. Molecular Immunology, 78, 113–120. 10.1016/J.MOLIMM.2016.09.002

Hernández-Martínez, P., Naseri, B., Navarro-Cerrillo, G., Escriche, B., Ferré, J., & Herrero, S. (2010). Increase in midgut microbiota load induces an apparent immune priming and increases tolerance to *Bacillus thuringiensis*. Environmental Microbiology, 12(10), 2730–2737. 10.1111/J.1462-2920.2010.02241.X

Horak, R. D., Leonard, S. P., & Moran, N. A. (2020). Symbionts shape host innate immunity in honeybees. Proceedings of the Royal Society B, 287(1933). 10.1098/RSPB.2020.1184

Jang, S., Ishigami, K., Mergaert, P., & Kikuchi, Y. (2024). Ingested soil bacteria breach gut epithelia and prime systemic immunity in an insect. Proceedings of the National Academy of Sciences of the United States of America, 121(11), e2315540121. 10.1073/PNAS.2315540121/SUPPL_FILE/PNAS.2315540121.SAPP.PDF

Johnston, P. R., & Rolff, J. (2013). Immune- and wound-dependent differential gene expression in an ancient insect. Developmental & Comparative Immunology, 40(3–4), 320–324. 10.1016/J.DCI.2013.01.012

Keshavarz, M., Franz, M., Xie, H., Zanchi, C., Mbedi, S., Sparmann, S., & Rolff, J. (2025). Immune-mediated indirect interaction between gut microbiota and bacterial pathogens. BMC Biology, 23(1), 1–18. 10.1186/S12915-025-02399-1/FIGURES/8

Korša, A., Baur, M., Schulz, N. K. E., Anaya-Rojas, J. M., Mellmann, A., & Kurtz, J. (2025). Experimental evolution of a pathogen confronted with innate immune memory increases variation in virulence. PLOS Pathogens, 21(6), e1012839. 10.1371/JOURNAL.PPAT.1012839

Korša, A., Lo, L. K., Gandhi, S., Bang, C., & Kurtz, J. (2022). Oral immune priming treatment alters microbiome composition in the red flour beetle *Tribolium castaneum*. Frontiers in Microbiology, 13, 793143. 10.3389/FMICB.2022.793143/TEXT

Lafont, M., Petton, B., Vergnes, A., Pauletto, M., Segarra, A., Gourbal, B., & Montagnani, C. (2017). Long-lasting antiviral innate immune priming in the Lophotrochozoan Pacific oyster, *Crassostrea gigas*. Scientific Reports 2017 7:1, 7(1), 1–14. 10.1038/s41598-017-13564-0

Lafont, M., Vergnes, A., Vidal-Dupiol, J., De Lorgeril, J., Gueguen, Y., Haffner, P., Petton, B., Chaparro, C., Barrachina, C., Destoumieux-Garzon, D., Mitta, G., Gourbal, B., & Montagnani, C. (2020). A sustained immune response supports long-term antiviral immune priming in the pacific oyster, *Crassostrea gigas*. MBio, 11(2). 10.1128/MBIO.02777-19/SUPPL_FILE/MBIO.02777-19-SF005.TIF

Lang, H., Duan, H., Wang, J., Zhang, W., Guo, J., Zhang, X., Hu, X., & Zheng, H. (2022). Specific Strains of Honeybee Gut *Lactobacillus* Stimulate host immune system to protect against pathogenic *Hafnia alvei*. Microbiology Spectrum, 10(1). 10.1128/SPECTRUM.01896-21/SUPPL_FILE/SPECTRUM01896-21_SUPP_1_SEQ8.PDF

Lanz-Mendoza, H., & Contreras-Garduño, J. (2022). Innate immune memory in invertebrates: Concept and potential mechanisms. Developmental and Comparative Immunology, 127. 10.1016/J.DCI.2021.104285

Li, M., Shao, D., Zhou, J., Gu, J., Qin, J., Chen, W., & Wei, W. (2020). Signatures within esophageal microbiota with progression of esophageal squamous cell carcinoma. Chinese Journal of Cancer Research = Chung-Kuo Yen Cheng Yen Chiu, 32(6), 755–767. 10.21147/J.ISSN.1000-9604.2020.06.09

Li, S., Nie, H., Huo, Z., & Yan, X. (2025). Transcriptomic signatures related to the immune priming of *Ruditapes philippinarum* in response to the re-infection of *Vibrio anguillarum*. Fish & Shellfish Immunology, 161, 110263. 10.1016/J.FSI.2025.110263

Li, Y., Song, X., Wang, W., Wang, L., Yi, Q., Jiang, S., Jia, Z., Du, X., Qiu, L., & Song, L. (2017). The hematopoiesis in gill and its role in the immune response of Pacific oyster *Crassostrea gigas* against secondary challenge with *Vibrio splendidus*. Developmental and Comparative Immunology, 71, 59–69. 10.1016/J.DCI.2017.01.024

Liu, C., Zhang, T., Wang, L., Wang, M., Wang, W., Jia, Z., Jiang, S., & Song, L. (2016). The modulation of extracellular superoxide dismutase in the specifically enhanced cellular immune response against secondary challenge of *Vibrio splendidus* in Pacific oyster (Crassostrea gigas). Developmental and Comparative Immunology, 63, 163–170. 10.1016/J.DCI.2016.06.002

Low, C. F., & Chong, C. M. (2020). Peculiarities of innate immune memory in crustaceans. Fish & Shellfish Immunology, 104, 605–612. 10.1016/J.FSI.2020.06.047

Martinez, C., Rodriguez, S., Vences, A., Barja, J. L., Toranzo, A. E., & Dubert, J. (2022). Role of the vibriolysin VemA secreted by the emergent pathogen *Vibrio europaeus* in the colonization of Manila clam mucus. Microorganisms, 10(12). 10.3390/MICROORGANISMS10122475

Mersni-Achour, R., Imbert-Auvray, N., Huet, V., Ben Cheikh, Y., Faury, N., Doghri, I., Rouatbi, S., Bordenave, S., Travers, M. A., Saulnier, D., & Fruitier-Arnaudin, I. (2014). First description of French V. tubiashii strains pathogenic to mollusk: II. Characterization of properties of the proteolytic fraction of extracellular products. Journal of Invertebrate Pathology, 123, 49–59. 10.1016/j.jip.2014.09.006

Mersni-Achour, R., Cheikh, Y. B., Pichereau, V., Doghri, I., Etien, C., Dégremont, L., Saulnier, D., Fruitier-Arnaudin, I., & Travers, M. A. (2015). Factors other than metalloprotease are required for full virulence of French *Vibrio tubiashii* isolates in oyster larvae. Microbiology, 161(Pt 5), 997–1007. 10.1099/mic.0.000058

Melillo, D., Marino, R., Italiani, P., & Boraschi, D. (2018). Innate immune memory in invertebrate metazoans: a critical appraisal. Frontiers in Immunology, 9, 372689. 10.3389/FIMMU.2018.01915/XML

Montagnani, C., Morga, B., Novoa, B., Gourbal, B., Saco, A., Rey-Campos, M., Bourhis, M., Riera, F., Vignal, E., Corporeau, C., Charrière, G. M., Travers, M. A., Dégremont, L., Gueguen, Y., Cosseau, C., & Figueras, A. (2024). Trained immunity: Perspectives for disease control strategy in marine mollusc aquaculture. Reviews in Aquaculture, 16(4), 1472–1498. 10.1111/RAQ.12906

Mottola, A., Bonerba, E., Figueras, M. J., Pérez-Cataluña, A., Marchetti, P., Serraino, A., Bozzo, G., Terio, V., Tantillo, G., & Di Pinto, A. (2016). Occurrence of potentially pathogenic arcobacters in shellfish. Food Microbiology, 57, 23–27. 10.1016/J.FM.2015.12.010

Naylor, R. L., Hardy, R. W., Buschmann, A. H., Bush, S. R., Cao, L., Klinger, D. H., Little, D. C., Lubchenco, J., Shumway, S. E., & Troell, M. (2021). A 20-year retrospective review of global aquaculture. Nature 2021 591:7851, 591(7851), 551–563. 10.1038/s41586-021-03308-6

Pradeu, T., & Du Pasquier, L. (2018). Immunological memory: What’s in a name? Immunological Reviews, 283(1), 7–20. 10.1111/IMR.12652

Prado, S., Romalde, J. L., Montes, J., & Barja, J. L. (2005). Pathogenic bacteria isolated from disease outbreaks in shellfish hatcheries. First description of *Vibrio neptunius* as an oyster pathogen. Diseases of Aquatic Organisms, 67(3), 209–215. 10.3354/dao067209

Prado, S., Dubert, J., & Barja, J. L. (2015). Characterization of pathogenic vibrios isolated from bivalve hatcheries in Galicia, NW Atlantic coast of Spain. Description of *Vibrio tubiashii* subsp. *europaeus* [corrected] subsp. nov. Systematic and Applied Microbiology, 38(1), 26–29. 10.1016/j.syapm.2014.11.005

Quast, C., Pruesse, E., Yilmaz, P., Gerken, J., Schweer, T., Yarza, P., Peplies, J., & Glöckner, F. O. (2012). The SILVA ribosomal RNA gene database project: improved data processing and web-based tools. Nucleic Acids Research, 41(Database issue), D590. 10.1093/NAR/GKS1219

Ramees, T. P., Dhama, K., Karthik, K., Rathore, R. S., Kumar, A., Saminathan, M., Tiwari, R., Malik, Y. S., & Singh, R. K. (2017). *Arcobacter*: an emerging food-borne zoonotic pathogen, its public health concerns and advances in diagnosis and control – a comprehensive review. Veterinary Quarterly, 37(1), 136–161. 10.1080/01652176.2017.1323355

Ramírez, C., Rojas, R., & Romero, J. (2020). Partial evaluation of autochthonous probiotic potential of the gut microbiota of *Seriola lalandi*. Probiotics and Antimicrobial Proteins, 12(2), 672–682. 10.1007/S12602-019-09550-9

Rey-Campos, M., Moreira, R., Gerdol, M., Pallavicini, A., Novoa, B., & Figueras, A. (2019). Immune tolerance in *Mytilus galloprovincialis* hemocytes after repeated contact with *Vibrio splendidus*. Frontiers in Immunology, 10(AUG), 471443. 10.3389/FIMMU.2019.01894/TEXT

Rey-Varela, D., Ana L. Diéguez, A.L., Rodriguez, S., Dinçtürk, E., Villada, A., Ojea, J., Polo, D., Dubert, J. (2025). Development of conventional and quantitative PCR assays for the detection of *Vibrio europaeus* using a species-specific gene identified from the pangenome. Aquaculture. (Manuscript submitted for publication)

Rodrigues, J., Brayner, F. A., Alves, L. C., Dixit, R., & Barillas-Mury, C. (2010). Hemocyte differentiation mediates innate immune memory in anopheles gambiae mosquitoes. Science, 329(5997), 1353–1355. 10.1126/SCIENCE.1190689/

Rojas, R., Blanco-Hortas, A., Kehlet-Delgado, H., Lema, A., Miranda, C. D., Romero, J., Martínez, P., Barja, J. L., & Dubert, J. (2021). First description outside Europe of the emergent pathogen Vibrio europaeus in shellfish aquaculture. Journal of invertebrate pathology, 180, 107542. 10.1016/j.jip.2021.107542

Saulnier, D., De Decker, S., Haffner, P., Cobret, L., Robert, M., & Garcia, C. (2010). A large-scale epidemiological study to identify bacteria pathogenic to Pacific oyster *Crassostrea gigas* and correlation between virulence and metalloprotease-like activity. Microbial ecology, 59(4), 787–798. 10.1007/s00248-009-9620-y

Smolowitz, R. (2024). Diseases of bivalves: historical and current perspectives. Diseases of Bivalves: Historical and Current Perspectives, 1–315. 10.1016/C2019-0-01464-1

Sompayrac, L. M. (2022). How the immune system works. John Wiley & Sons.

Turner, M., Van Hulzen, L., Guse, K., Agany, D., & Pietri, J. E. (2024). The gut microbiota confers resistance against *Salmonella Typhimurium* in cockroaches by modulating innate immunity. IScience, 27(12), 111293. 10.1016/J.ISCI.2024.111293/

Travers, M. A., Mersni Achour, R., Haffner, P., Tourbiez, D., Cassone, A. L., Morga, B., Doghri, I., Garcia, C., Renault, T., Fruitier-Arnaudin, I., & Saulnier, D. (2014). First description of French V. tubiashii strains pathogenic to mollusk: I. Characterization of isolates and detection during mortality events. Journal of Invertebrate Pathology, 123, 38–48. 10.1016/j.jip.2014.04.009

Wang, H., Yang, B., Li, Q., & Liu, S. (2024). Low-dose of formalin-inactivated *Vibrio alginolyticus* protects *Crassostrea gigas* from secondary infection and confers broad-spectrum *Vibrio* resistance on offspring. Developmental and Comparative Immunology, 152. 10.1016/J.DCI.2023.105122

Wang, J., Wang, L., Yang, C., Jiang, Q., Zhang, H., Yue, F., Huang, M., Sun, Z., & Song, L. (2013). The response of mRNA expression upon secondary challenge with *Vibrio anguillarum* suggests the involvement of C-lectins in the immune priming of scallop *Chlamys farreri*. Developmental & Comparative Immunology, 40(2), 142–147. 10.1016/J.DCI.2013.02.003

Yue, F., Zhou, Z., Wang, L., Ma, Z., Wang, J., Wang, M., Zhang, H., & Song, L. (2013). Maternal transfer of immunity in scallop *Chlamys farreri* and its trans-generational immune protection to offspring against bacterial challenge. Developmental & Comparative Immunology, 41(4), 569–577. 10.1016/J.DCI.2013.07.001

Zannella, C., Mosca, F., Mariani, F., Franci, G., Folliero, V., Galdiero, M., Tiscar, P. G., & Galdiero, M. (2017). Microbial diseases of bivalve mollusks: infections, immunology and antimicrobial defense. Marine Drugs 2017, Vol. 15, Page 182, 15(6), 182. 10.3390/MD15060182

Zhang, T., Qiu, L., Sun, Z., Wang, L., Zhou, Z., Liu, R., Yue, F., Sun, R., & Song, L. (2014). The specifically enhanced cellular immune responses in Pacific oyster (*Crassostrea gigas*) against secondary challenge with Vibrio splendidus. Developmental & Comparative Immunology, 45(1), 141–150. 10.1016/J.DCI.2014.02.015

Zhang, X., Guo, M., Sun, Y., Wang, Y., & Zhang, Z. (2022). Transcriptomic analysis and discovery of genes involving in enhanced immune protection of Pacific abalone (*Haliotis discus hannai*) in response to the re-infection of *Vibrio parahaemolyticus*. Fish & Shellfish Immunology, 125, 128–140. 10.1016/J.FSI.2022.04.045

