## Supplemental Table S1-S3 Suplemental Figures S1-S6 for "Oral immune priming modulates microbiota composition and supports pathogen control in the Manila clam (*Ruditapes philippinarum*)"

### *Supplementary Material*

#### 1 Supplementary Tables

For more information on Supplementary Material and for details on the different file types accepted, please see here.

**Table S1. Primers, probe and DNA standard used for the *ktr*A-based qPCR assay (Rey-Varela et al., 2025).**

| **Primers** | **Sequence (5’→3’)** | **Amplicon size (bp)** |
|---|---|---|
| Veuro-F | GGCACACTTTACAGTAATTG | 148 |
| Veuro-R | ACAAATCACAGCTTGTGTAAG | |
| Veuro-probe | /6-FAM/ACA CTT AGG /ZEN/GCA TTC GGT GAC AGG G/IABkFQ/ | |
| Standard DNA | ATGGCACACTTTACAGTAATTGGGCTAGGACGATTTGGCATAGCCGCAAGTTTAGAGCTCATACACTTAGGGCATTCGGTGACAGGGGTAG ATCGAGAGTCGAAAATAGTGAGTAAGTATGTTGAAGCGCTTACACAAGCTGTGATTTGTGA | 152 |

**Table S2. Statistical results of data processing.**

Clean reads are the raw sequences filtered the low quality, that is Clean reads which are finally used for subsequent analysis; Clean data is the number of bases in the final Clean reads; AvgLen is the average length of Clean reads; Q20 and Q30 are the percentage of bases that have quality values in Clean reads greater than 20 (sequencing error rate less than 1%) and 30 (sequencing error rate less than 0.1%); GC (%) represents the content of GC bases in Clean reads; Effective (%) represents the percentage of the number of Clean reads to the number of Raw PE.

| **Sample** | **Clean_reads** | **Base** | **AvgLen** |
|---|---|---|---|
| P1 | 22614 | 32744766 | 1447 |
| P2 | 23200 | 33540529 | 1445 |
| P3 | 21201 | 30475226 | 1437 |
| P4 | 22609 | 32610772 | 1442 |
| P5 | 21826 | 31743489 | 1454 |
| P6 | 21293 | 30800763 | 1446 |
| P7 | 22261 | 32396513 | 1455 |
| P8 | 23694 | 34397135 | 1451 |
| P9 | 23636 | 34352986 | 1453 |
| P10 | 21703 | 30991847 | 1427 |
| P11 | 22450 | 32487463 | 1447 |
| NP0.5 | 23475 | 34207232 | 1457 |
| NP1 | 21259 | 30687223 | 1443 |
| NP1.5 | 21939 | 32035426 | 1460 |
| NP2 | 23534 | 34283665 | 1456 |
| NP2.5 | 22916 | 33511970 | 1462 |
| P+SC0.5 | 22299 | 32365409 | 1451 |
| P+SC1 | 23423 | 34352670 | 1466 |
| P+SC1.5 | 23407 | 33795423 | 1443 |
| P+SC2 | 22663 | 32854556 | 1449 |
| P+SC2.5 | 21862 | 31799107 | 1454 |
| P+SC5.5 | 23903 | 34412263 | 1439 |

**Table S3. Collapsed ASVs by Eztaxon.**

| Collapsed ASV | Species | Original ASVs |
|---|---|---|
| ASVB1 | *Streptococcus thermophilus/S. salivarius/S. vestibularis* (*Streptococcus*) | ASV33, ASV36, ASV40, ASV95, ASV106, ASV143, ASV139, ASV133 |
| ASVB2 | *Vibrio gigantis/V. celticus/V. coralliirubri/V. artabrorum/V. bathopelagicus/V. crassostreae/V. pomeroyi (Vibrio)* | ASV44, ASV56, ASV81, ASV76, ASV124, ASV149, ASV147 |
| ASVB3 | *Psychrilyobacter atlanticus/P. piezotolerans (Psychrilyobacter)* | ASV18, ASV22, ASV49, ASV53, ASV52, ASV105, ASV104 |
| ASVB4 | *Vibrio europaeus* | ASV0, ASV2, ASV7, ASV8, ASV9 |
| ASVB5 | FM995181_s (Bacteria) | ASV5, ASV14, ASV17, ASV21, ASV25 |
| ASVB6 | *Escherichia fergusonii/Shigella flexneri/E. coli/S. sonnei (Escherichia)* | ASV89, ASV80, ASV68, ASV134 |
| ASVB7 | *Acinetobacter lwoffii/Prolinoborus fasciculus/A. pecorum (Acinetobacter)* | ASV3, ASV11, ASV16, ASV6 |
| ASVB8 | *Poseidonibacter lekithochrous* | ASV15, ASV27, ASV42 |
| ASVB9 | *Cutibacterium acnes* | ASV39, ASV47, ASV120 |

| | | |
|---|---|---|
| ASVB10 | *Pseudoalteromonas distincta/P. elyakovii/P. atlantica/P. paragorgicola/P. arctica/P. agarivorans/P. espejiana/P. carrageenovora/P. undina/P. nigrifaciens/P. hodoensis (Pseudoalteromonas)* | ASV127, ASV115, ASV82 |
| ASVB11 | *Anaplasmataceae* | ASV24, ASV28 |
| ASVB12 | *Sphingobium limneticum* | ASV41, ASV45 |
| ASVB13 | *Brevundimonas huaxiensis/B. vesicularis/B. nasdae/B. fontaquae/B. intermedia/B. huaxiensis (Brevundimonas)* | ASV20, ASV26 |
| ASVB14 | *Algibacter* | ASV135, ASV145 |
| ASVB15 | CP014505_s (*Caballeronia*) | ASV34, ASV117 |

#### 2 Supplementary Figures

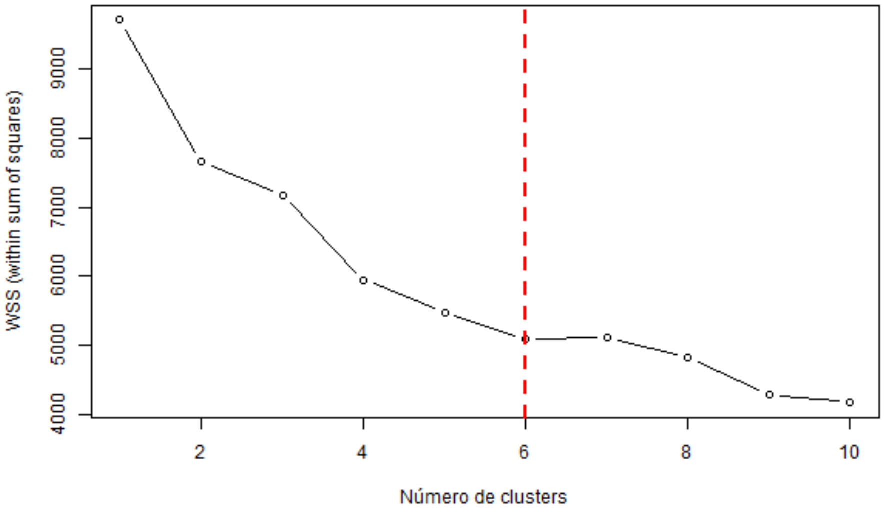


**Supplementary Figure S1. Determination of the optimal number of clusters using the Elbow Method.** The within-cluster sum of squares (WSS) was calculated for different numbers of clusters (k). The inflection point at $k = 6$ indicates the optimal number of clusters used for subsequent analyses.

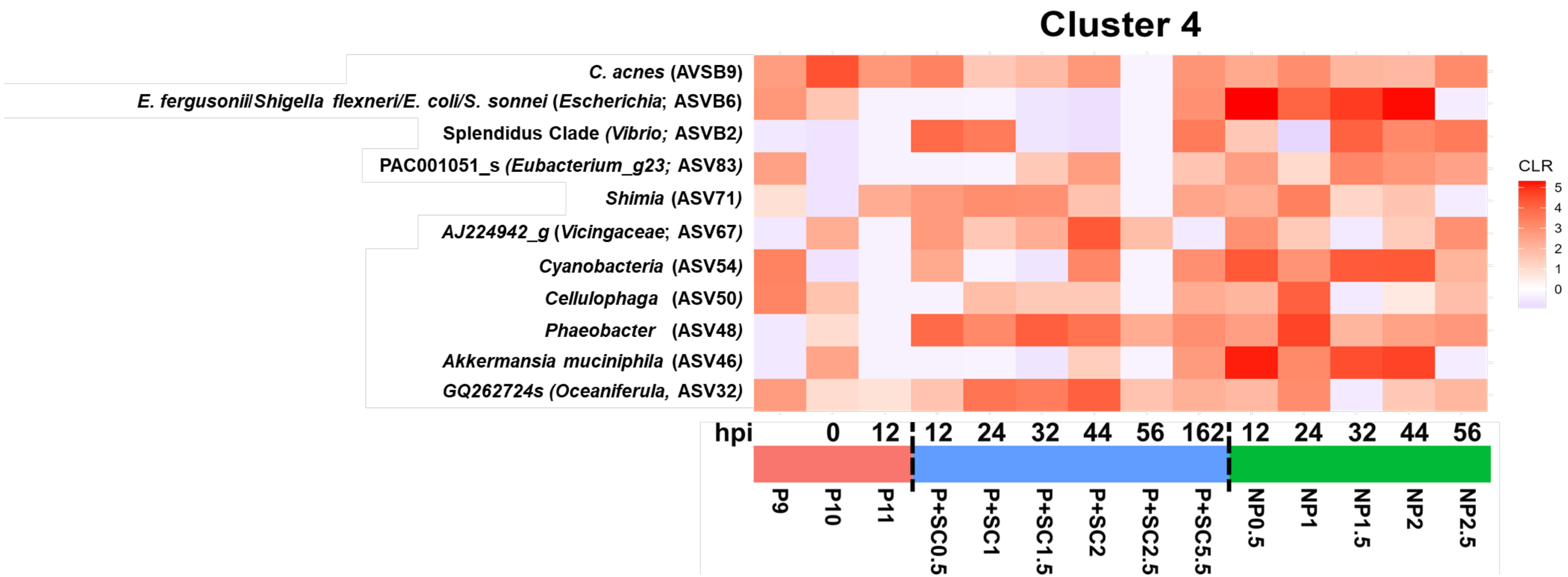


**Supplementary Figure S2. Cluster 4.**

Characteristics are showed on Fig. 5.

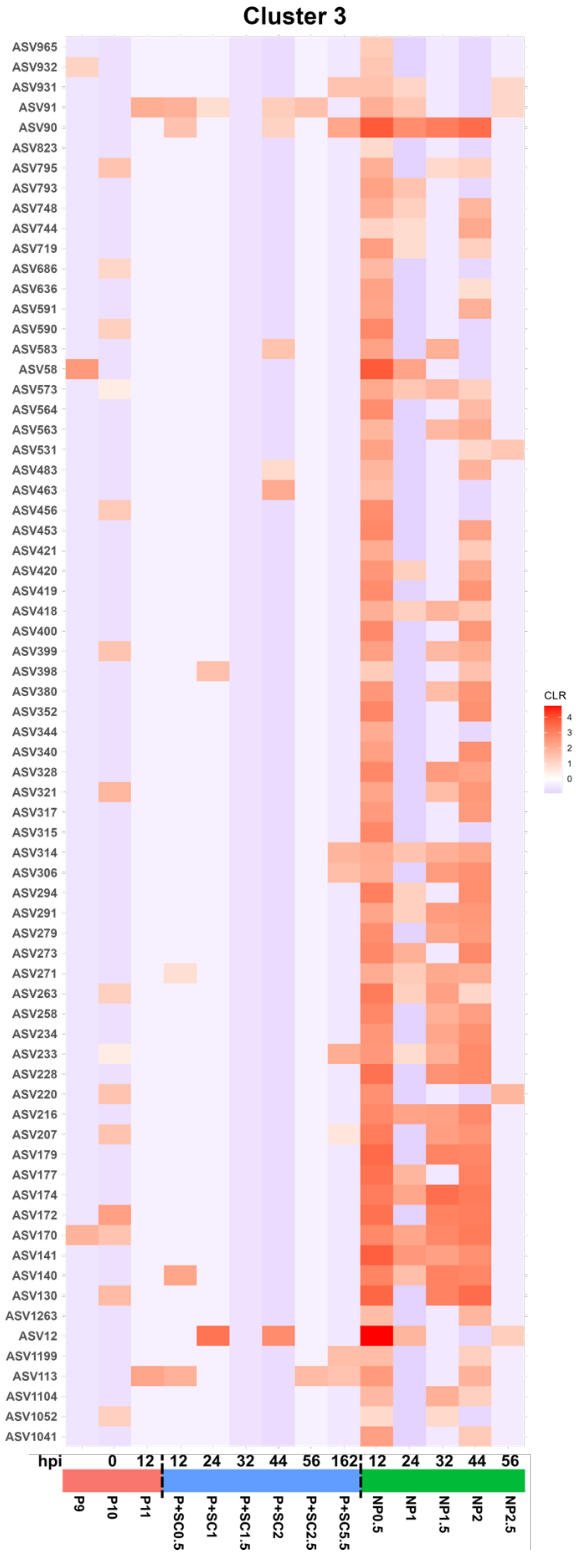


**Supplementary Figure S3. Cluster 3.**

Characteristics are showed on Fig. 5.

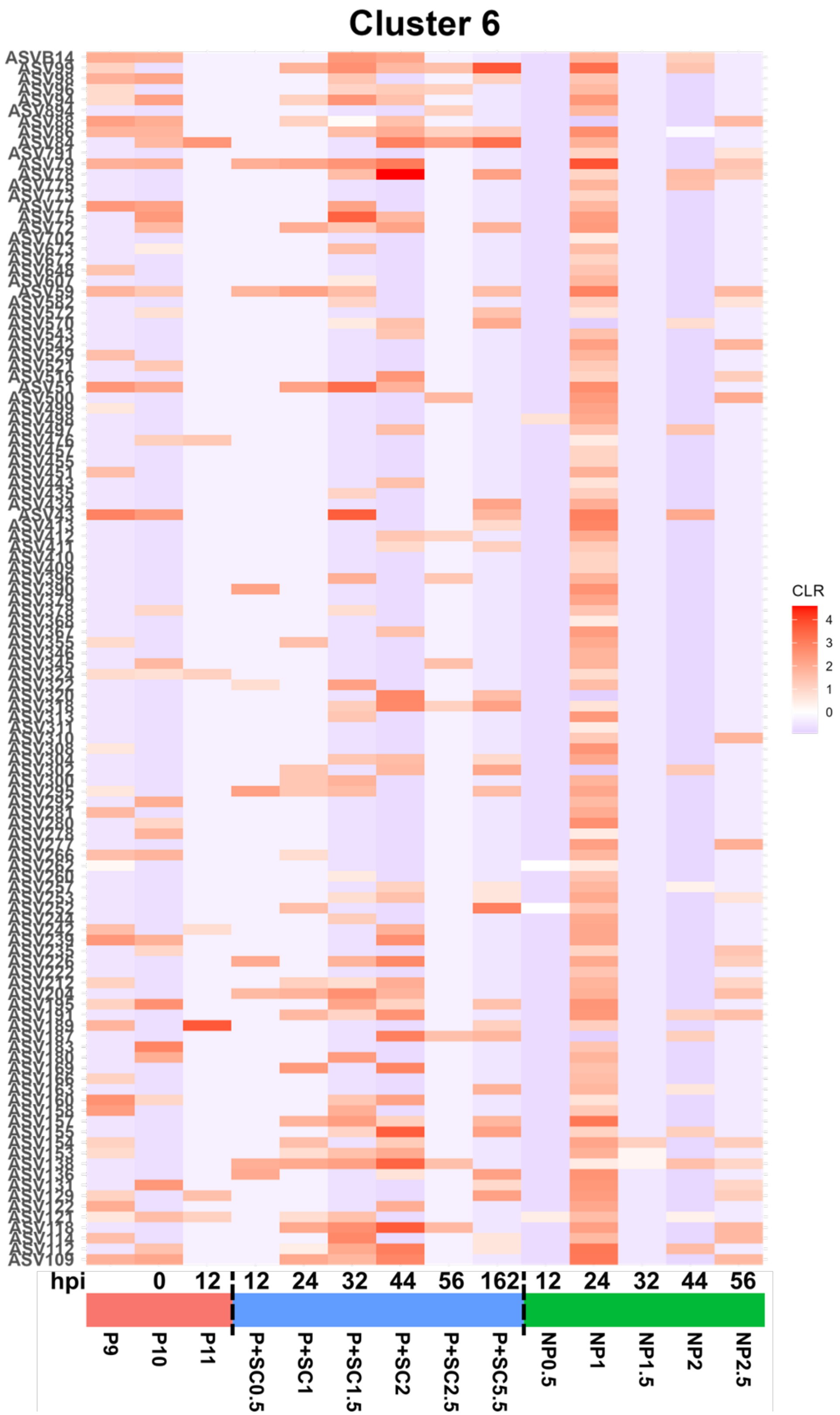


**Supplementary Figure S4. Cluster 6.**

Characteristics are showed on Fig. 5

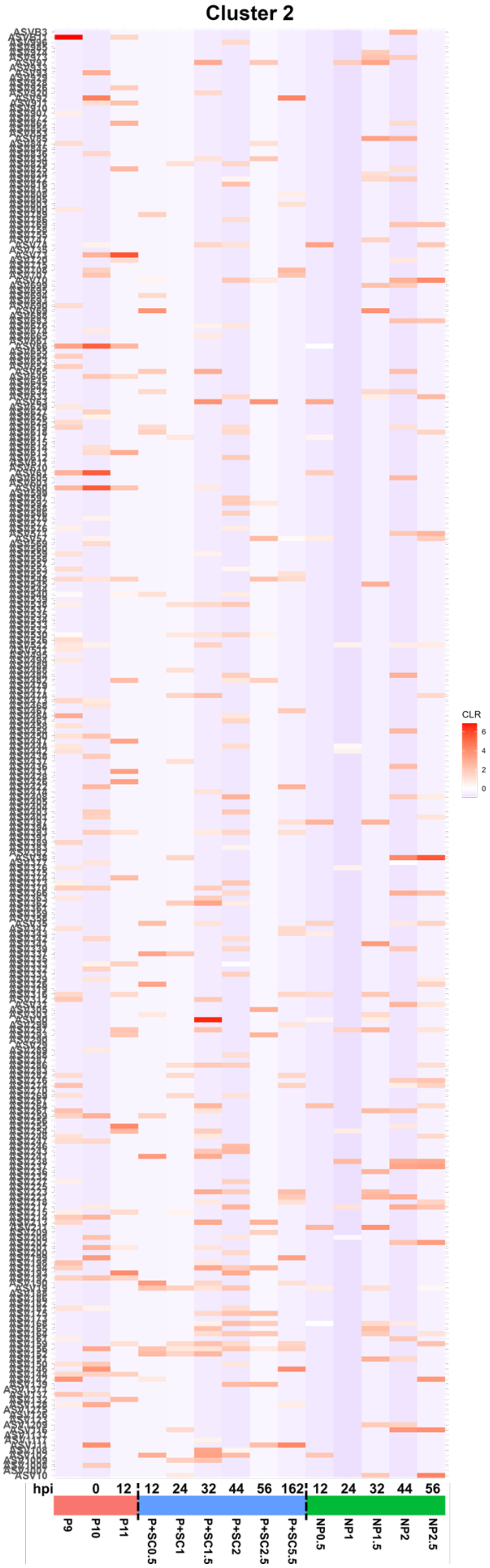


**Supplementary Figure S5. Cluster 2.**

Characteristics are showed on Fig. 5
